# A self-organized geometry sensing oscillator spatially regulates cell division in archaea

**DOI:** 10.64898/2026.07.23.738820

**Authors:** Wenchao Zheng, Shan Zhao, Yafei Liu, Zhanyi Chen, Jiarong Wu, Xiaoyao Zou, Jiaoyao Cui, Xiangdong Chen, Joe Lutkenhaus, Fabai Wu, Shishen Du

**Affiliations:** State Key Laboratory of Metabolism and Regulation in Complex Organisms, College of Life Sciences, Wuhan University, Wuhan, Hubei, China; Hubei Key Laboratory of Cell Homeostasis, College of Life Sciences, Wuhan University, Wuhan, Hubei, China; School of Biomedical Engineering, College of Engineering, Eastern Institute of Technology, Ningbo, China; State Key Laboratory of Virology and Biosafety, College of Life Sciences, Wuhan University, Wuhan, Hubei, China; Department of Microbiology, Molecular Genetics and Immunology, University of Kansas Medical Center, Kansas City, Kansas, USA

## Abstract

Cell division must be tightly regulate in time and space. Many bacteria and eukaryotes employ geometry sensing systems to regulate cell division spatiotemporally. However, how archaea ensure they divide in the right place at the right time is little understood. Here, we report the discovery of a geometry sensing oscillator that determines the division plane in the model haloarcheon *Haloferax volcanii*, which relies on two tubulin-like proteins, FtsZ1 and FtsZ2, for division. This <u>arc</u>haeal <u>o</u>scillator (named as Arco system) is composed of three components, ArcoA, ArcoB and ArcoC, which are a membrane associated Ras superfamily GTPase, a SepF-like protein and an antagonist of FtsZ1, respectively. ArcoA interacts with both ArcoB and ArcoC. ArcoAB sense the geometry of the cell to oscillate between the two cell poles, generating a time-averaged ArcoC gradient which is high at the cell poles but low at the midcell, where FtsZs can assemble into the Z ring to initiate cytokinesis. As a result, deletion of the Arco system caused aberrant FtsZ assembly throughout the cell, leading to irregular division and formation of minicells. Interestingly, the Arco system is globally distributed among archaea and co-occurs with FtsZ1, suggesting that it emerged in the last archaeal common ancestor (LACA). The oscillatory behavior and phenotypes of the archaeal Arco system is remarkably analogous to the bacterial Min system and yet share no protein homology, representing a remarkable example of convergent evolution. Overall, these results suggest that archaea, similar to bacteria and eukaryotes, independently evolved self-organized Turing reaction diffusion systems to sense geometry and spatially regulate cell division.

## Introduction

The ability to undergo autonomous division is a hallmark of cellular life. To achieve proper distribution of genetic material and other cellular content into daughter cells, the division machineries must be well positioned in both prokaryotic and eukaryotic cells. One prominent way to establish a cell division site precisely is to integrate cell geometry to orchestrate a spatial gradient of proteins that negatively regulate division proteins (Fig. 1a). In the fission yeast *Schizosaccharomyces pombe,* Pom1 proteins form a gradient decreasing from cell poles to the cell center, thereby constraining Cdr2 proteins to establish a cortical node and recruit downstream proteins to form a contractile ring at midcell ^1–3^. Analogously, the Min system of the rod-shaped *Escherichia coli* form a dynamic oscillation between the two cell poles, forming a time-averaged protein gradient to prevent the tubulin-like FtsZ protein from polymerizing into Z ring (scaffold of the division apparatus) at the two poles ^4–6^. *Bacillus subtilis* employs a diversified Min system persistently anchored to the cell poles to regulate Z ring positioning, while *Caulobacter crescentus* uses a system consisted of MipZ, ParB and PopZ to integrate status of chromosome segregation and cell curvature to restrict Z ring formation to roughly the midcell ^7^.

**Fig. 1.**
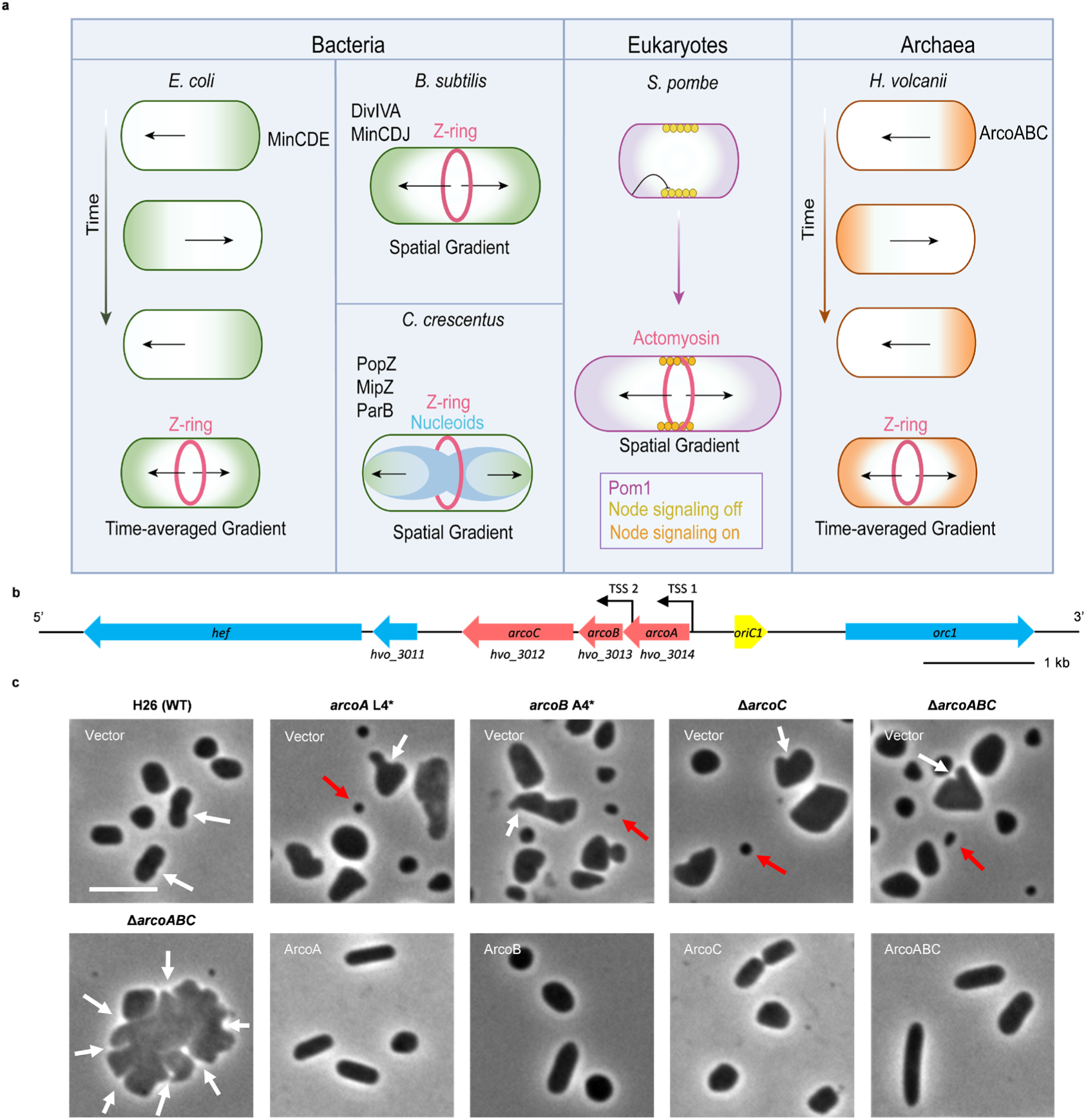
Spatial regulation of cell division cross life and identification of the Arco system in *H. volcanii*. a. Convergent evolution of geometry-sensing systems involved in division site placement across life. Min system in bacteria: In *E. coli*, MinD/MinE oscillate pole-to-pole to create a MinC gradient to permit Z-ring assembly exclusively at the cell center. In *B. subtilis*, MinC/D are persistently anchored to the cell poles by DivIVA/MinJ (no oscillation) so that Z ring assembly is prevented at the cell poles but not in the midcell. MipZ gradient in *C. cresentus*: the FtsZ inhibitor MipZ forms a gradually decreasing gradient on the nucleoid from the cell pole toward the cell center by interacting with ParB, which is anchored to the cell poles by PopZ. As a result, Z ring formation is allowed in the midcell but not at the cell poles. Pom1 gradient in fission yeast: Pom1 forms a polar gradient that delays mitotic entry until cell length reaches a threshold, thereby regulates positioning of the Actomyosin ring. Arco system in archaea: ArcoA/ArcoB oscillate from pole-to-pole to create a bipolar ArcoC gradient that gradually decreases from the cell poles to midcell, restricting Z-ring assembly exclusively at the cell center. All these systems achieve geometry-dependent division plane placement by creating a time-average gradient of division inhibitors that is high at the cell poles but low at the cell center, however, the proteins involved are completely unrelated. b. Genomic organization of the *arco* operon (*arcoA*, *arcoB*, *arcoC*) near the replication origin *oriC1*. Two transcription start sites (TSS) are indicated by black arrows. Scale bar, 1 kb. c. Representative images of wild type H26, *arcoA* L4*, *arcoB* A4*, Δ*arcoC*, and *arco* operon deletion cells, with or without plasmid-borne complementation. Minicells are indicated by red arrows, and surface indentations, which represent the division sites, are indicated by white arrows. For complementation, the corresponding gene was expressed from its native promoter on a plasmid. Scale bar, 5 µm.

While Pom1 gradient indirectly rely on microtubules for polar recruitment, the Min oscillation patterns are self-organized without the help of cytoskeleton ^3^. Instead, they orchestrate via a simple-yet-elegant Turing reaction-diffusion mechanism ^8–11^, primarily achieved by 1) the ATP-dependent cooperative binding of MinD to the membrane, 2) the ATPase stimulation and subsequent membrane-unbinding by MinE, and 3) the free diffusion of unbound MinD and MinE in the cytosol ^12^. MinD proteins then carry MinC as a cargo, which depolymerizes FtsZ at undesired locations such as the poles ^4^. The overarching roles of these gradient-forming systems are to sense the size and shape of a cell, i.e. geometry sensing, and transform the geometrical information into a positional information for the cell division machinery ^3^.

Although several gradient-forming mechanisms have been reported to regulate cell division spatiotemporally in bacteria and eukaryotes, none have been found in archaea. Many archaeal lineages employ FtsZ to scaffold the assembly of the division apparatus (divisome), a mechanism thought to already existed in the last archaeal common ancestor (LACA) ^13–15^. Many archaea such as the haloarchaeon *Haloferax volcanii* encode two essential FtsZ paralogs, FtsZ1 and FtsZ2 ^16^. Formation of the Z ring in *H. volcanii* relies on CdpC and SepF, which serve as membrane anchor for FtsZ1 and FtsZ2, respectively ^13,17,18^. Once the Z ring is assembled, SepF recruits downstream division proteins such as CdpA and CdpBs to form the complete divisome ^19–21^. Although distant homologs of MinD ATPases also exist in haloarchaea, they were found to regulate motility and have no association with FtsZ-based cell division ^22,23^; no other gradient forming system has been reported so far in archaea. Hence, the mechanism underlying FtsZ-utilizing archaea to position their cell division machineries has been long sought-after.

In this study, we report an <u>arc</u>haeal <u>o</u>scillator (Arco) that positions the Z ring in *H. volcanii*, with homologs existing broadly across the archaea domain (Fig. 1a). This tripartite system is composed of a membrane-binding GTPase (ArcoA), a regulator of the GTPase (ArcoB), and an antagonist of FtsZ1 (ArcoC), each functionally analogous to the MinD, MinE, and MinC of the bacterial Min system and yet share no evolutionary relation with the latter (Fig. 1a). Archaea thus independently evolved a Turing reaction-diffusion pattern for the spatial positioning of cell division.

## Results

### The Arco system is important for cell division and morphology

To identify potential spatial regulators of FtsZs in *H. volcanii*, we carried out *in vivo* crosslinking coupled with immunoprecipitation and mass spectrometry (CLIP-MS) screening using GFP fusions of FtsZ1 and FtsZ2 as the baits and GFP as a control. Proteins with a >8-fold enrichment over the control and a P-value <0.05 were considered as potential FtsZ1 or FtsZ2 interactors. 158 proteins were enriched in the FtsZ1-GFP immunoprecipates, including 4 previously known cell division proteins and 154 uncharacterized proteins; while 65 proteins were enriched in the FtsZ2-GFP immunoprecipates, with 8 known division proteins and 57 unstudied proteins (Fig. S1a-b, Table S1-2). Among these, 38 proteins were enriched in both the FtsZ1-GFP and FtsZ2-GFP immunoprecipitates, including 4 known division proteins or candidate division proteins, CdpA, CdpC, HVO_1130 and HVO_1512 (a SepF-like protein) (Fig. S1c, Table S3).

To determine if the remaining enriched proteins are involved in cell division or function as regulators, we examined their subcellular distribution using GFP fusions. Most of the proteins were evenly distributed in the cytoplasm or formed bipolar punctates (Table S4). Among them, we noticed HVO_3012 (ArcoC), which was previously reported to lead to vesicle formation upon deletion ^24^, localized to midcell and form polar caps in a subset of cells (Fig. S1d). Previous studies have shown that *hvo_3012* and its two upstream genes, *hvo_3013* (*arcoB*) and *hvo_3014 (arcoA)*, form an operon called *oap*, as it is adjacent to the replication origin *oriC1* ^25^ (Fig. 1b). Interestingly, similar to HVO_3012, GFP-HVO_3013 also localized to midcell in a subpopulation of cells and it also localized asymmetrically in some cells, however, GFP-HVO_3014 showed an asymmetric distribution toward the poles or sides (Fig. S1d). Structural and functional prediction of the Arco proteins by AlphaFold 3 ^26^ and the HHpred ^27^ algorithm suggested that ArcoA is a Ras superfamily GTPase with a unique N-terminal amphipathic helix, while ArcoB is a SepF-like protein with residues 90-98 likely forming an amphipathic helix (Fig. S1e-j; Table S5). ArcoC is a hypothetical protein with no known homologs; it is predicted to contain an N-terminal zinc-ribbon domain and a C-terminal segment, which are connected by a long intrinsically disordered linker (residues 35-280).

To test if the Arco proteins participate in cell division, we generated single-gene knockout strains as well as a full operon deletion strain. Strikingly, deletion of any individual *arco* gene or the entire operon all caused morphological abnormalities and division defects (Fig. 1c and Fig. S2a). Instead of the typical rod or disc shape, the mutant cells were irregular, accompanied by the appearance of abundant minicells. Moreover, many mutant cells appeared to undergo asymmetric division (Fig. 1c and Fig. S2a). Complementation of the deletion strains with the respective gene or the full operon from a plasmid showed that *arcoC* and the full operon could complement in trans, but expression of ArcoA or ArcoB could not rescue the division and morphological defects of the corresponding deletion strain (Fig. 1c and Fig. S2a). This was likely due to polar effects of *arcoA* or *arcoB* deletion on the transcription of downstream genes since previous studies showed that this operon has two transcription start sites (TSS), one upstream of *arcoA* and another within *arcoA*, which together generate five transcripts (Fig. 1b) ^25^. To avoid the polar effects, we generated *arcoA* and *arcoB* null strains (*arcoA* L4*, *arcoB* A4*) by introducing stop codons at the fourth codon of the respective gene in the genome. Similar to the Δ*arcoA* and Δ*arcoB* cells, *arcoA* L4* and *arcoB* A4* cells displayed morphological abnormalities and division defects (Fig. 1c). However, unlike the full-frame deletion strains, plasmid-borne expression of ArcoA or ArcoB fully rescued the defects of the *arcoA* L4* and *arcoB* A4* cells. Hence, using premature stop codons in *arcoA* and *arcoB* avoids polar effects, providing a more rigorous and reliable approach to study their functions. These results thus demonstrate that the Arco system is important for the prevention of polar cell division which leads to minicells, akin to the Min system in *E. coli*.

Next, we explored the respective roles of Arco proteins by increasing their expression levels. To do this, we overexpressed individual Arco proteins or the entire *arco* operon using a xylose-inducible promoter ^28^ in wild type cells. Interestingly, overexpression of ArcoA or ArcoC caused cell enlargement and irregular division, whereas overexpression of ArcoB led to the generation of elongated cells and minicells (Fig. S2b). Notably, ArcoA was extremely toxic since it resulted in morphological defects even without xylose induction, presumably due to leaky expression from the xylose promoter. Intriguingly, overexpression of the full *arco* operon resulted in elongated cells and minicells, similar to overexpression of ArcoB (Fig. S2b), suggesting that the relative stoichiometry of the Arco proteins, rather than their absolute levels, is critical for their proper function. In summary, these results suggest that ArcoA and ArcoC may be more directly involved in cell division inhibition, while ArcoB may play a regulatory role.

### The Arco system regulates cell division site placement in *H. volcanii*

To investigate whether the Arco system affect cell division and morphology by acting on division proteins, we examined the localization of FtsZs and other key division proteins in the absence of Arco proteins or when they were overexpressed. Strikingly, in the absence of ArcoA, ArcoC, or the entire system, all the tested division proteins assembled into disorganized fibers and spirals throughout the cell, instead of the typical band-like Z ring structure (Fig. 2a and Fig. S3a). Although the absence of ArcoB also led to mislocalization of the division proteins, much fewer fibers were observed (Fig. 2a and Fig. S3a). Interestingly, overexpression of ArcoA or ArcoC in wild-type cells resulted in the formation of FtsZ1-/FtsZ2-GFP foci or aggregates, implying that they block FtsZ1-/FtsZ2-GFP assembly. By contrast, overexpression of ArcoB caused both FtsZ1-GFP and FtsZ2-GFP to form disorganized filamentous structures throughout the cells (Fig. 2b and Fig. S3b), resembling the phenotype observed in the absence of ArcoA or ArcoC. These results again suggest that ArcoA and ArcoC work together to inhibit Z-ring assembly, whereas ArcoB counteracts their function and restrict FtsZ assembly to the midcell.

**Fig. 2.**
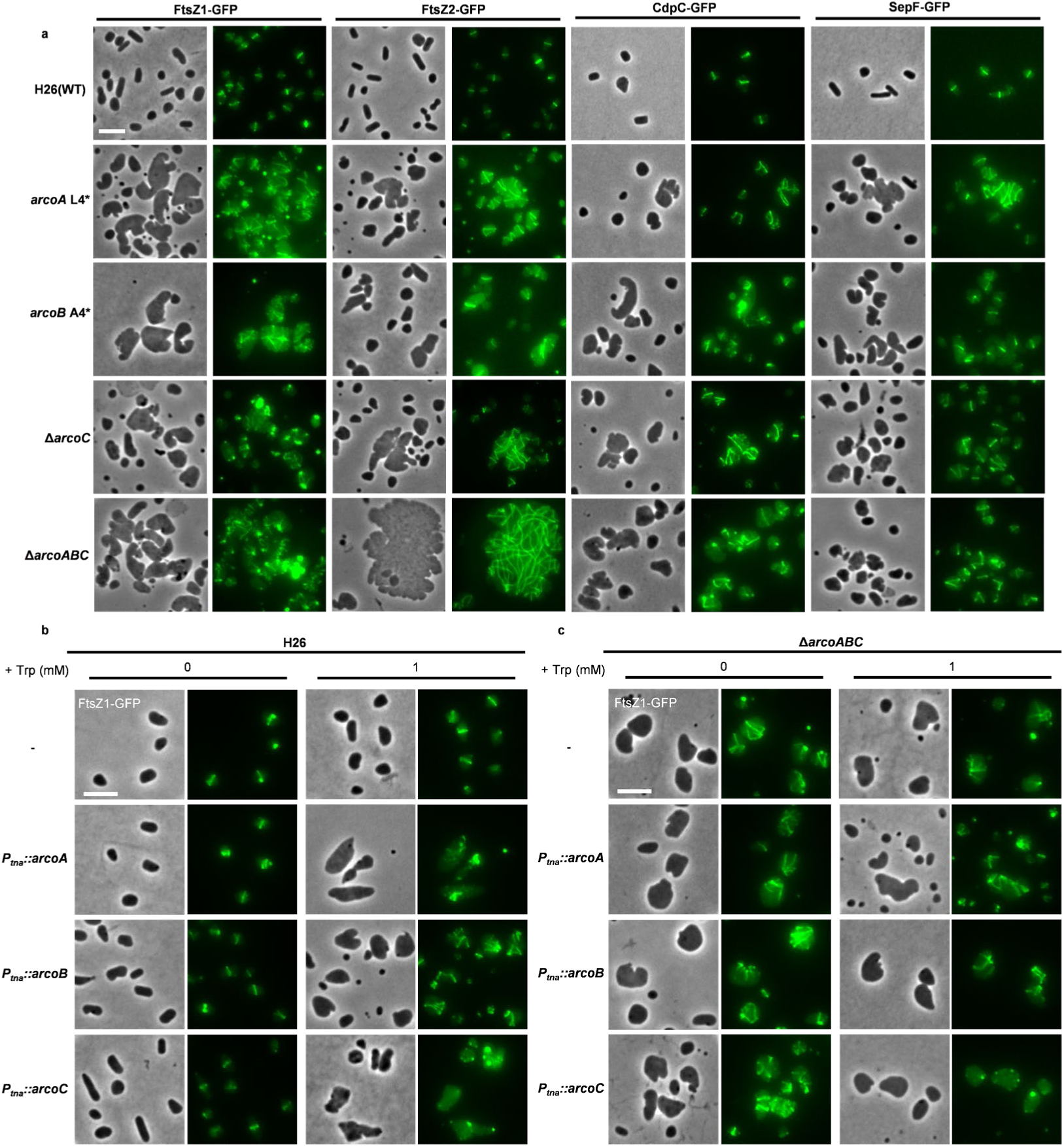
The Arco system controls localization of divisome proteins. a. Representative images of the localization of GFP fusions of FtsZ1, FtsZ2 and their membrane anchors CdpC, SepF in wild type H26, *arcoA* L4*, *arcoB* A4*, Δ*arcoC*, and *arco* operon deletion strains. All GFP fusion proteins were expressed under the control of their native promoters from plasmids, respectively. b. Representative images of the FtsZ1-GFP localization upon overexpression of ArcoA, ArcoB or ArcoC in wild type H26 backgrounds. FtsZ1-GFP (controlled by its native promoter) and ArcoA, ArcoB or ArcoC (controlled by *P_tna_* promoter) were co-expressed from plasmids. Cells were grown at 45 °C overnight with the indicated concentrations of tryptophan (Trp). c. Representative images of the FtsZ1-GFP localization upon overexpression of ArcoA, ArcoB or ArcoC in *arco* operon deletion backgrounds. FtsZ1-GFP (controlled by its native promoter) and ArcoA, ArcoB or ArcoC (controlled by *P_tna_* promoter) were expressed from plasmids. Cells were grown at 45 °C overnight with the indicated concentrations of tryptophan (Trp). Scale bars, 5 μm.

To test if the mis-localization of division proteins upon Arco protein overexpression depended on the presence of the endogenous Arco system, we overexpressed them in the Δ*arcoABC* cells. Interestingly, overexpression of ArcoA or ArcoB no longer affected the localization of FtsZ1-/FtsZ2-GFP which formed mis-organized fibers within these irregular cells. However, overexpression of ArcoC in the Δ*arcoABC* cells prevented FtsZ1-/FtsZ2-GFP from forming fibers (Fig. 2c, Fig. S3c). These observations indicate that ArcoC alone can inhibit Z-ring assembly while ArcoA and ArcoB rely on the presence of ArcoC to exert their effects. Overall, we conclude that the Arco system regulates Z-ring assembly and thereby controls the site of cell division.

### The Arco system is a geometry sensing oscillator that regulates Z ring positioning

To further examine how Arco system spatially regulates cell division proteins, we performed time-lapse imaging of GFP-tagged Arco proteins in wild type H26 cells. Strikingly, both ArcoA-GFP and ArcoB-GFP underwent pole-to-pole oscillation in rod-shaped cells (Fig. 3a, b and Supplementary Video 1-2). In disc-shaped cells, the oscillation mode switched to a circumferential movement along the cell periphery (Fig. S4a, Supplementary Video 3). ArcoC-GFP also oscillated, albeit with a slightly different pattern: the cap-like structures at the cell poles alternately and periodically increased and decreased in intensity (Fig. 3c, Supplementary Video 4). These results confirm that the Arco proteins constitute a geometry sensing oscillator, reminiscent of the bacterial Min system.

**Fig. 3.**
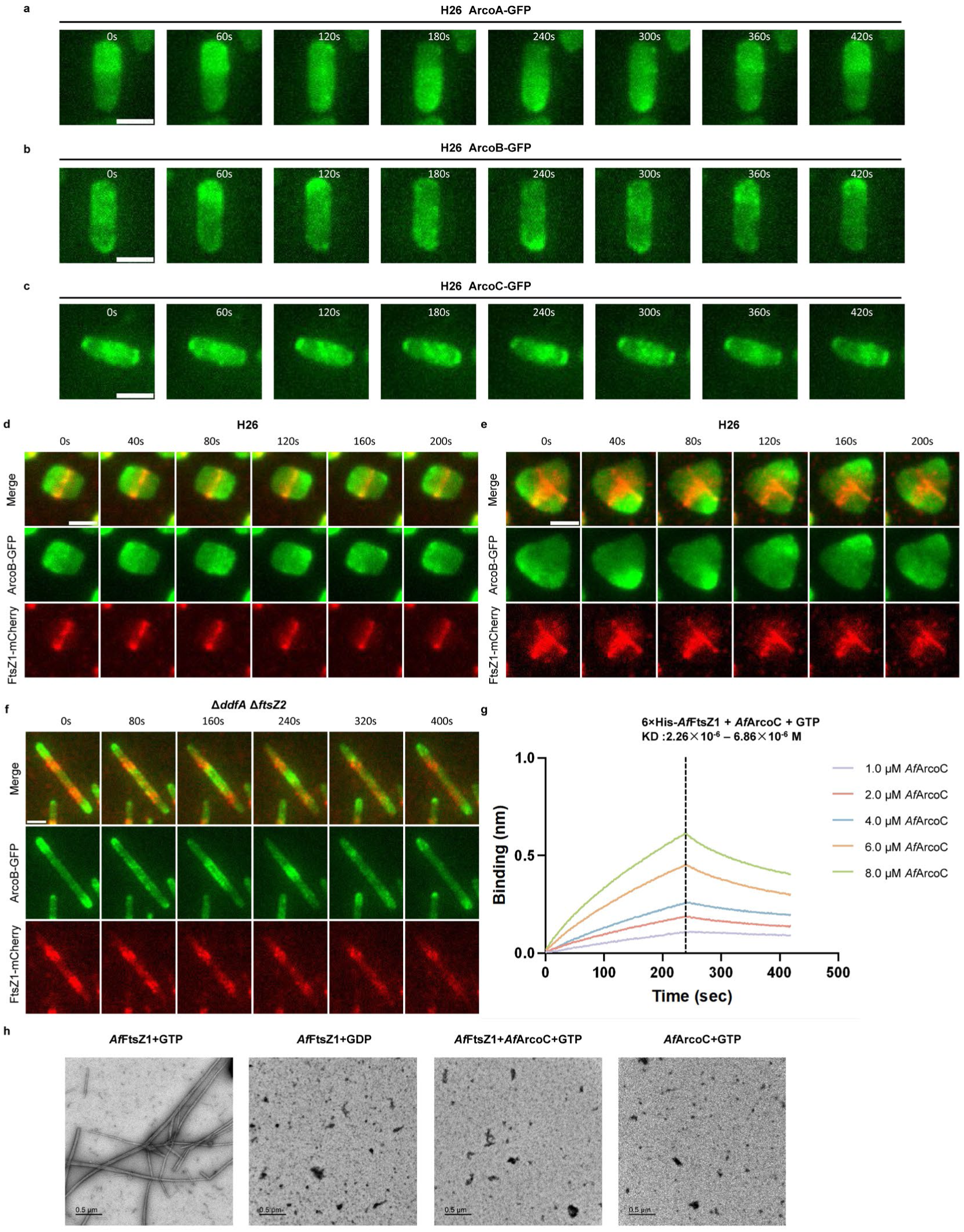
Arco proteins oscillate and control Z ring positioning. a-c. Representative time-lapse images of ArcoA-GFP (a), ArcoB-GFP (b) and GFP-ArcoC (c) in rod-shaped H26 cells. ArcoA-GFP was expressed from a plasmid under the control of the *P_tna_* promoter (induced with 1 mM Trp). ArcoB-GFP was expressed from a plasmid under the control of the *P_xyl_* promoter (induced with 0.5 mM xylose). GFP-ArcoC was expressed from a plasmid driven by the native promoter of the *arco* operon. All strains were grown at 45 °C overnight before imaging. Time in seconds. Scale bar, 2 µm. d-e. Representative time-lapse images of FtsZ1-mCherry and ArcoB-GFP in rod-shaped and triangular-shaped wild type H26 cells. FtsZ1-mCherry was expressed from its native promoter and ArcoB-GFP was under the control of the *P_xyl_* promoter on the same plasmid. Cells were grown at 45 °C overnight with 0.5 mM xylose. Time in seconds. Scale bar, 2 µm. f. Representative time-lapse images of FtsZ1-mCherry and ArcoB-GFP in elongated rod-shaped Δ*ddfA* Δ*ftsZ2* cells. Samples were prepared as in e and f. Scale bar, 2 µm. g. BLI assay showing that *Af*ArcoC interacts with *Af*FtsZ1 *in vitro*. Details about the experimental set up and procedures are described in Methods. The concentration of the 6×his-*Af*FtsZ1 protein remains constant (0.5 μM), the concentration of GTP is 1 mM, while the concentration of *Af*ArcoC varies. h. Representative images of transmission electron microscopy (TEM) of *Af*FtsZ1 incubated with or without *Af*ArcoC. The polymerization reactions were set up in a 50 μL volume as described in Methods. The final concentration of *Af*FtsZ1 and *Af*ArcoC was 2.5 μM, GTP/GDP was 1mM. Scale bar, 0.5 µm.

To investigate the relationship between the Arco oscillator and Z-ring placement, we examined the localization dynamics of ArcoB-GFP and FtsZ1-mCherry in wild type H26 cells by time-lapse imaging. In rod-shaped cells, ArcoB-GFP oscillated along the long axis between the two poles, while FtsZ1-mCherry formed a ring persistently at the midpoint (Fig. 3d, Supplementary Video 5). In triangular-shaped cells, ArcoB-GFP oscillated sequentially among the three poles, and FtsZ1-mCherry fibers accumulated between each pair of poles, converging at the cell center (Fig. 3e, Supplementary Video 6). To determine whether Z-ring positioning was determined by the Arco oscillation or it just assembled in the cell’s geometric center regardless of the Arco oscillation, we examined ArcoB-GFP and FtsZ1-mCherry localization dynamics in elongated rod-shaped cells. To this end, we constructed a Δ*ddfA*Δ*ftsZ2* strain: deletion of *ddfA* locks the cells in rod shape ^29^, while deletion of *ftsZ2* causes cell elongation ^16^. Strikingly, in these rod-shaped elongated cells, ArcoB-GFP no longer oscillated simply from pole to pole. Instead, two oscillations were observed, each of which oscillated from the pole to the cell center (Fig. 3f, Supplementary Video 7). Accordingly, FtsZ1-mCherry accumulated at the midpoints of the two oscillations (roughly the one-quarter and three-quarter along the cell length), rather than forming a single ring at midcell. These results strongly indicate that the Arco oscillator determines the site of Z-ring formation. Also, since the Arco oscillator can restrict FtsZ1-ring positioning in the absence of FtsZ2, it likely acts on FtsZ1 rather than FtsZ2.

To confirm that the Arco system acts through FtsZ1 instead of FtsZ2, we examined ArcoB-GFP oscillation and FtsZ1 or FtsZ2 localization in strains where FtsZ2 or FtsZ1 expression was under the control of a tryptophan-inducible *P_tna_* promoter ^16^. When FtsZ2 was depleted (by removing tryptophan from the medium), multiple oscillation foci appeared in the large aberrant cells, yet FtsZ1-mCherry still correctly localized to the center of each oscillation (Fig. S5a, Supplementary Video 8). In contrast, upon FtsZ1 depletion, ArcoB-GFP continued to oscillate, but FtsZ2-mCherry were dispersed throughout the cells (Fig. S5b, Supplementary Video 9). Therefore, the Arco system targets FtsZ1 rather than FtsZ2 to regulate cell division.

### ArcoC blocks Z ring formation by inhibiting FtsZ1 polymerization

Given that overexpression of ArcoC inhibited Z-ring formation and the Arco systems likely acts on FtsZ1, we hypothesized that ArcoC interacts with FtsZ1 and inhibits its assembly. To test this, we examined the interaction between ArcoC and FtsZ1 using the split-FP assay (based on the principle that interaction between two proteins of interest can reconstitute superfolder GFP ^30^) in the Δ*arcoABC* background. We observed fibrous localization and detected strong fluorescence signals when ArcoC and FtsZ1 fused to complementary GFP fragments were co-expressed (Fig. S6a, b). In contrast, no fluorescence was detected when only one of the fusions or neither was expressed. We similarly observed fibrous localization and strong fluorescence signals when ArcoC and FtsZ2 fused to complementary GFP fragments were co-expressed (Fig. S6c-d). However, Co-IP experiments showed that FtsZ1-Flag, but not FtsZ2-Flag, co-immunoprecipitated with ArcoC-GFP (Fig. S6e-h), suggesting that ArcoC interacts directly with FtsZ1 but not FtsZ2 *in vivo*.

To further confirm the interaction between FtsZ1 and ArcoC, we tried to purify *H. volcanii* ArcoA and ArcoB and tested their interaction *in vitro*. However, the attempts failed, so we purified homologues of ArcoC and FtsZ1 from *Archaeoglobus fulgidus* and tested their interaction by Biolayer Interferometry (BLI) experiments. As shown in Fig. 3g, *Af*ArcoC robustly bound to 6×His*-Af*FtsZ1 immobilized on the biosensor, with a dissociation constant (KD) ranging from 1.33×10^-7^ to 3.15×10^-6^ M, demonstrating that they directly interact with each other. Following this, we examined the effect of *Af*ArcoC on *Af*FtsZ1 polymerization by transmission electron microscopy. *Af*FtsZ1 assembled into filamentous structures after incubation with GTP but not GDP (Fig. 3h), consistent with FtsZ’s requirement of GTP to polymerize ^31^. Strikingly, addition of *Af*ArcoC to the reaction completely eliminated the *Af*FtsZ1 filaments, indicating that *Af*ArcoC antagonizes *Af*FtsZ1 polymerization (Fig. 3h). Taken together, these results demonstrate that ArcoC interacts directly with FtsZ1 and inhibits its polymerization, but the oscillation restricts its action to the poles.

### ArcoA and ArcoB are necessary and sufficient for oscillation while AroC oscillates with them by interacting with ArcoA

To investigate the mechanism underlying the oscillation, we examined the dynamics of Arco proteins in single *arco* gene null strains and the *arco* operon deletion strain. In the absence of ArcoA, oscillations of both ArcoB and ArcoC were abolished (Fig. S4b). Similarly, the absence of ArcoB eliminated the oscillation of ArcoA and ArcoC. By contrast, deletion of *arcoC* did not affect oscillations of ArcoA or ArcoB (Fig. S4b). These results suggest that ArcoA and ArcoB are necessary and sufficient for the oscillation, whereas ArcoC is not necessary. In line with this, co-expression of ArcoA and ArcoB in the Δ*arcoABC* cells reconstituted their oscillations (Fig. 4a-b), suggesting that they interact with each other to generate the oscillation. In support of this, Split-FP assays showed that ArcoA and ArcoB interacted with each other in the absence of ArcoC (in Δ*arcoABC* background) (Fig. S7a-b). Moreover, Co-IP experiments showed that ArcoA-HA and ArcoB-GFP co-precipitates with each other (Fig. 4c, Fig. S7c). Subsequently, we tried to confirm their interaction *in vitro*. However, attempts to purify *H. volcanii* ArcoA and ArcoB failed. Therefore, we purified their homologues from *A. fulgidus* and examined their interaction by BLI assays. As shown in Fig. 4d, *Af*ArcoB rapidly bound to 6×His*-Af*ArcoA immobilized on the biosensor and also dissociated quickly, with a dissociation constant (KD) ranging from 2.73×10^-7^ to 6.78×10^-7^ M. Based on these results, we concluded that ArcoA and ArcoB constitute the oscillation within the cell, analogous to *E. coli* MinD and MinE, while ArcoC is a passenger of the ArcoAB oscillator (analogous to MinC), presumably by interacting with ArcoA.

**Fig. 4.**
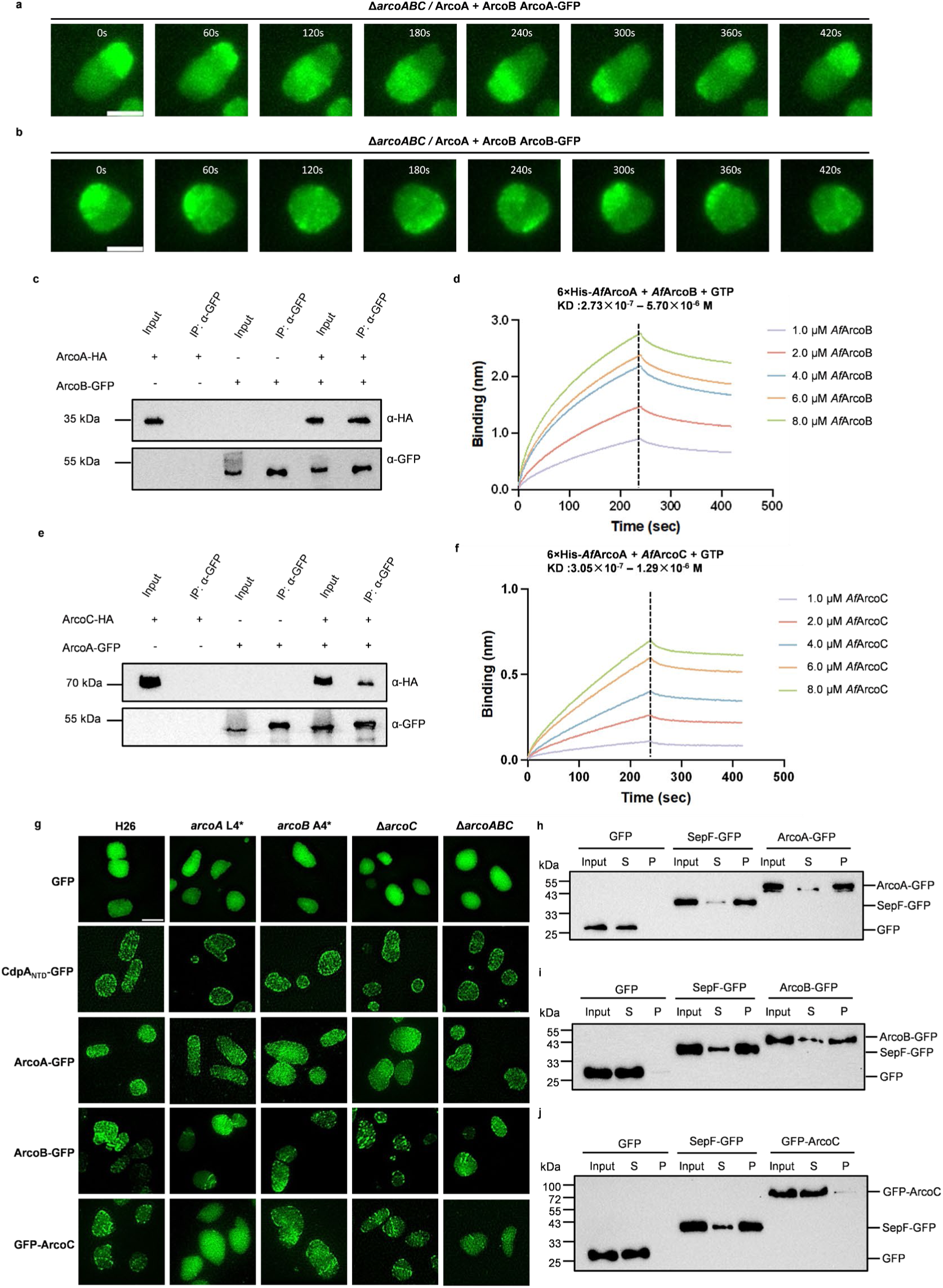
Investigations of the mechanism underlying the Arco oscillation. a-b. Representative time-lapse images of the ArcoAB oscillation in Δ*arcoABC* cells. A single plasmid harboring untagged *arcoA* and *arcoB* under the control of the native promoter of the *arco* operon, and *arcoA-gfp* (a) or *arcoB-gfp* (b) under the control of the *P_tna_* promoter was employed for this experiment. Cells were grown at 45 °C overnight and expression of ArcoA-GFP or ArcoB-GFP was induced with 1 mM Trp. Time in seconds. Scale bars, 2 µm. c. Co-IP experiment showing that ArcoA interacts with ArcoB *in vivo*. Cultures of *H. volcanii* expressing the indicated proteins were lysed by sonication; supernatants were incubated with GFP antibodies coated magnetic beads. Immunocomplexes were eluted with boiling SDS-PAGE loading buffer and then analyzed by immunoblot. d. BLI assay showing that *Af*ArcoA interacts with *Af*ArcoB *in vitro*. Details about the experimental set up and procedures are described in Methods. The concentration of the 6×his-*Af*ArcoA protein remains constant (0.5 μM), the concentration of GTP is 1 mM, while the concentration of *Af*ArcoB varies. e. Co-IP experiment showing that ArcoC interacts with ArcoA *in vivo*. Cultures of *H. volcanii* expressing the indicated proteins were lysed by sonication; supernatants were incubated with GFP antibodies coated magnetic beads. Immunocomplexes were eluted with boiling SDS-PAGE loading buffer and then analyzed by immunoblot. f. BLI assay showing that *Af*ArcoC interacts with *Af*ArcoA *in vitro*. Details about the experimental set up and procedures are described in Methods. The concentration of the 6×his-*Af*ArcoA protein remains constant (0.5 μM), the concentration of GTP is 1 mM, while the concentration of *Af*ArcoC varies. g. Representative 2D-SIM images of the localization of GFP-tagged Arco proteins in indicated genetic backgrounds. ArcoA-GFP was expressed from a plasmid under the control of the *P_tna_* promoter (induced with 1 mM Trp). ArcoB-GFP was expressed from a plasmid under the control of the *P_xyl_* promoter (induced with 0.5 mM xylose). GFP-ArcoC was expressed from a plasmid driven by the native promoter of the *arco* operon. All strains were grown at 45 °C overnight before imaging. Scale bar, 2 μm. h-j. b. Membrane fractionation assays to determine the location of the Arco proteins *in vivo*. Cultures of *arco* operon deletion strains expressing the indicated proteins were lysed by sonication; the membrane fraction was isolated by ultracentrifugation; the pellet was resuspended to the original volume and analyzed along with the supernatant by immunoblot. (h) ArcoA-GFP, (i) ArcoB-GFP, (j) ArcoC-GFP, SepF-GFP was employed as a positive control. S, supernatant (cytosol); P, pellet (membrane).

To test if there is an interaction between ArcoA and ArcoC *in vivo*, we again employed the split-FP assays in the Δ*arcoABC* background. When ArcoA and ArcoC fused to complementary GFP fragments were co-expressed, we observed fluorescent foci and some fibrous structures, accompanied by strong fluorescence signals (Fig. S7d, e). However, no fluorescence was detected when only one of the fusions or neither was expressed. Consistently, Co-IP experiments showed that ArcoC-HA and ArcoA-GFP co-immunoprecipitated (Fig. 4e, Fig. S7f). Furthermore, purified ArcoA and ArcoC of *A. fulgidus* interacted robustly, with a dissociation constant (KD) ranging from 1.20×10^-7^ to 1.29×10^-6^ M (Fig. 4f). Taken together, these results demonstrate that ArcoC interacts with ArcoA and it likely oscillates with the ArcoAB oscillator via its interaction with ArcoA.

### Membrane binding of ArcoA and ArcoB is important for oscillation

Both MinD and MinE of the bacterial Min oscillator must bind the membrane to generate the oscillation ^12,32,33^. By analogy, we suspected that both ArcoA and ArcoB should associate with the membrane. In support of this, structural predictions revealed that both of them contain an amphipathic helix (Fig. S1e-g, S8a-b and S9a-b). To confirm that they are associated with the membrane, we first examined the localization of their GFP fusions by Structured Illumination Microscopy (SIM). In wild type H26 cells, as well as in the single *arco* gene null strains and the Δ*arcoABC* strain, ArcoA-GFP and ArcoB-GFP exhibited a membrane localization pattern, similar to that of the transmembrane helix of CdpA (CdpA_NTD_) (Fig. 4g). In contrast, ArcoC-GFP showed membrane localization in wild-type, *arcoB* A4* and Δ*arcoC* cells, however, it was uniformly distributed throughout the cytoplasm in *arcoA* L4* or Δ*arcoABC* cells (Fig. 4g). Membrane fractionation assays showed that ArcoA and ArcoB were largely enriched in the membrane fraction (pellet), similar to the membrane associated protein SepF (Fig. 4h, i). By contrast, ArcoC was almost exclusively present in the supernatant (cytosol) (Fig. 4j). Taken together, these results demonstrate that ArcoA and ArcoB can bind the membrane independently, whereas ArcoC likely associates with the membrane indirectly through binding ArcoA.

To test if the amphipathic helices of ArcoA and ArcoB were critical for membrane binding, we compared the localization of GFP-tagged ArcoA, ArcoB and their variants, including ArcoA_Δ1-20_ (full-length ArcoA lacking the amphipathic helix), ArcoA_1-20_ (the amphipathic helix alone), 2×ArcoA_1-20_ (two copies of the amphipathic helix connected by a linker), ArcoB_Δ89-97_ (full-length ArcoB lacking the amphipathic helix), ArcoB_89-97_ (the amphipathic helix alone), and 2×ArcoB_89-97_ (two copies of the helix connected by a linker). Note that the expression level of ArcoA_1-20_-GFP and ArcoB_89-97_-GFP were undetectable (Fig. S8c, S9c), so that subsequent experiments were performed using the tandem constructs. Using 2D-SIM, we observed that ArcoA_Δ1-20_ and ArcoB_Δ89-97_ showed a uniform cytosolic distribution, whereas the tandem helix constructs 2×ArcoA_1-20_ and 2×ArcoB_89-97_ independently localized to the membrane (Fig. S8d, S9d). In agreement with this, membrane fractionation assays confirmed that 2×ArcoA_1-20_-GFP and 2×ArcoB_89-97_-GFP were largely enriched in the membrane (pellet), whereas ArcoA_Δ1-20_-GFP and ArcoB_Δ89-97_-GFP were primarily found in the cytosol (supernatant) (Fig. S8e, S9e), confirming that the amphipathic helices mediate membrane binding of ArcoA and ArcoB.

To test if membrane binding is important for ArcoAB function, we performed complementation test and examined the oscillation. As expected, ArcoA_Δ1-20_-GFP and ArcoB_Δ89-97_-GFP were unable to rescue the morphological and division defects of the *arcoA* L4* *and arcoB* A4* strains (Fig. S8f, S9f). Moreover, they also failed to oscillate in cells (Fig. S8g, S9g, Supplementary Video 10-11). Altogether, these results demonstrate that the amphipathic helices of ArcoA and ArcoB mediate their association with the membrane, which is critical for oscillation and regulation of cell division.

### The Arco system is widely conserved among Archaea and co-occurs with FtsZ

Establishing the Arco system as a spatial regulator of cell division in *H. volcanii*, we examined its distribution in archaea. Mapping the *arcoABC* gene cluster onto the global archaeal phylogeny revealed a broad distribution across the major clades of extant archaea, including most of the best sampled Euryarchaeota superphylum except for Poseidoniia, Thermococci, and Hadarchaeota (Fig. 5a). ArcoABC operons are also present in about half of DPANN archaeal phyla, although they are known for sub-micron sizes. Among Asgard archaea, it exists in Jordiarchaeia as full operons, while in Heimdallarchaeia, *arcoC* is mostly located apart from the *arcoAB* gene pair although full arcoABC operons existed in 8% of the genomes examined (Fig. 5a). Finally, no Arco systems were found in genomes of the TACK superphylum, coinciding with the general lack of FtsZ1 in these clades. Such a distribution suggested that Arco systems are widespread and highly correlated with FtsZ1-based cell division systems.

**Figure 5.**
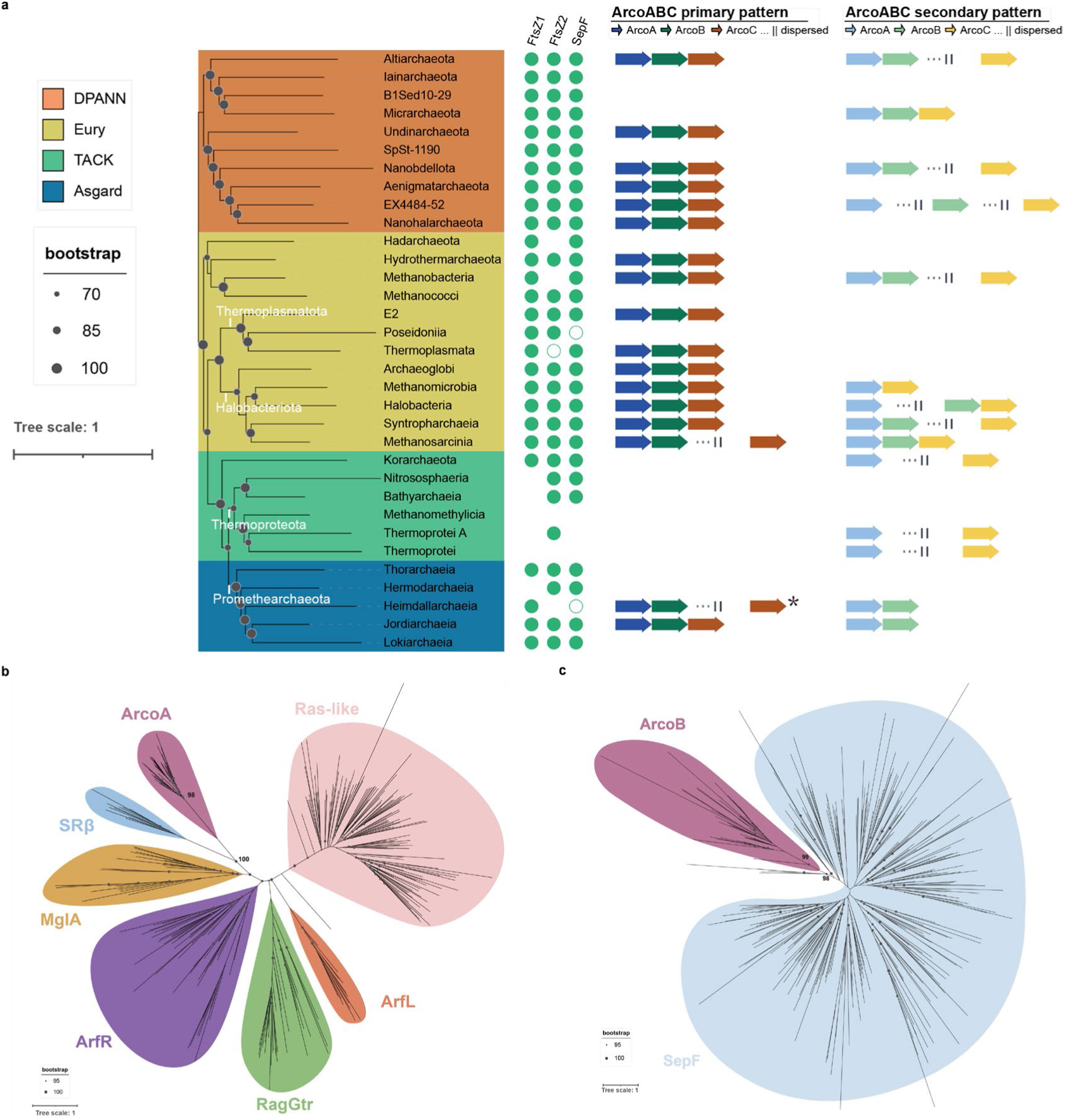
Global distribution and evolutionary origin of the Arco system. a, Distribution of the *arcoABC* gene clusters in genomically well-sampled lineages across the domain of Archaea. The phylogenetic tree was constructed using a representative species from each lineage using a concatenated archaeal protein marker set and LG+F+G4 model. The presence of FtsZ1, FtsZ2 and SepF homologues is indicated by circles beside each lineage; open circles denote detection in 10–50% of genomes within the lineage, whereas filled circles denote detection in more than 50% of genomes. The primary and secondary gene arrangements observed within each lineage are indicated in the middle and right. The asterisk indicates that a subset of Heimdallarchaeia also contains a continued arcoABC operon. b-c, Phylogenetic trees of ArcoA and ArcoB homologs, respectively, analyzed using LG+F+C60+G4 model with 2000 ultrafast bootstraps. Bootstrap values above 95 are indicated.

Next, we set out to examine the evolutionary origins of the components of the Arco system. Maximum likelihood analyses of global Ras GTPase superfamily suggested that ArcoA are more related to the Asgard-eukaryotic Arf family, while distant from the Asgard-eukaryotic Ras-like family (Fig. 5b). Notably, ArcoA forms a monophyletic clade sister to the SRβ family proteins with high bootstrap support. SRβ has a single N-terminal transmembrane helix and play an essential role in the co-translational targeting of secretory and membrane proteins to the endoplasmic reticulum ^34^. It is highly conserved across eukaryotes, but its prokaryotic origin has been unknown. We found that the membrane-binding N-terminal extension is highly conserved across the ArcoA family proteins, unlike the remaining cytosolic GTPases including the basal prokaryotic MglA family. It is thus plausible that the eukaryotic SRβ is evolutionarily related to archaeal ArcoA.

Maximum likelihood analyses suggested that ArcoA-associated ArcoB proteins form a monophyletic clade sister to the SepF proteins (Fig. 5c). SepF is distributed widely across bacteria and archaea and has been suggested to be involved in the cell division of LECA ^13,15^. The wide distributions of archaeal ArcoB and their monophyly outside SepF suggested that it may have originated from an ancestral gene duplication after the bacteria-archaea split.

In summary, the overall distribution of Arco systems and monophyly of ArcoA and ArcoB suggested that they likely have emerged in the last archaeal common ancestor (LACA) to regulate FtsZ1-based cell division and have been lost in specific lineages during early archaeal diversification.

## Discussion

In this study, we demonstrate that the Arco system operates as a geometry sensing oscillator that determines the position of the Z-ring by directly inhibiting FtsZ1 polymerization in *H. volcanii* (Fig. 1a). The Arco system co-occurs with FtsZ1 in archaea and resembles the bacterial Min system, but it employs completely unrelated proteins. This suggests that archaea and bacteria independently evolve geometry sensing systems to regulate FtsZ-based cell division since their ancestral split. Thus, this study not only provides critical insights into the spatial regulation of archaeal cell division but also reveals that geometry sensing regulates cell division in all three domains of life. Also, the discovery of the Arco system opens the avenue for reconstitution of the archaeal cell division machinery for synthetic cell ^35,36^.

The Arco system consists of the ArcoABC proteins. Among them, ArcoA (Ras superfamily GTPase) and ArcoB (SepF-like) form the oscillator and ArcoC is a passenger of the oscillation by binding to ArcoA. We showed that both ArcoA and ArcoB bind to the membrane through an amphipathic helix and membrane binding is critical for generation of the oscillation and regulation of cell division. ArcoC is a potent inhibitor of FtsZ1, it blocks FtsZ1 assembly both *in vivo* and *in vitro*. Based on our results, we propose the following model for regulation of Z ring positioning by the Arco system. ArcoA (GTP-bound) and ArcoB generate the pole-to-pole oscillation through a mechanism analogous to the MinD-MinE system in bacteria. ArcoC binds to ArcoA and is carried along with the ArcoAB oscillator. ArcoC (in complex with ArcoA) directly binds FtsZ1 and prevents its polymerization on the membrane, thereby blocking Z-ring assembly throughout the cells. However, the ArcoAB oscillation creates a time-averaged ArcoC concentration gradient that is high at the cell poles but low at the cell center, allowing FtsZ1 assembles into the Z ring at midcell but not the poles (Fig. 6). Once the Z ring position is established, other division proteins such FtsZ2 and CdpBs are recruited sequentially to assemble into the complete divisome. This model explains why deletion of *arcoA*, *arcoC* or the entire operon leads to disorganized FtsZ1 fibers throughout the cell (inappropriate Z-ring initiation at non-midcell sites), and why overexpression of ArcoC suppresses FtsZ1 polymerization and causes cell enlargement. The antagonistic role of ArcoB – which counteracts ArcoA/ArcoC – is consistent with ArcoB stimulating the GTPase activity of ArcoA, thereby releasing the complex from the membrane and sustaining the oscillation. One of the important role of the Arco system is to achieve geometry sensing, namely transforming the information of cell size and shape into positional cues for cell division; this represents one of the most fundamental biophysical principles to regulate cellular life ^3^. In triangular cells, ArcoB cycles among three poles and Z-rings form between each pair; in elongated cells with two oscillation, two Z-rings appear at the midpoints of each oscillation, but not at the geometric center. Thus, the oscillation pattern itself is the readout of cell shape and size, dictating division site placement. The bacterial Min systems have been shown to achieve geometry sensing by aligning with symmetric axis and adapting its length scale to the cellular boundary, properties that are well encoded within its innate reaction-diffusion network ^37,38^. The Arco systems are also encoded in archaea with a broad range of cell shapes including sub-micron-sized DPANN archaea and tentacled Asgard archaea^14^. It would thus be exciting to quantitatively explore the nature and mechanism of geometry sensing for the Arco system across phyla and morphologies in the domain of archaea.

**Fig. 6.**
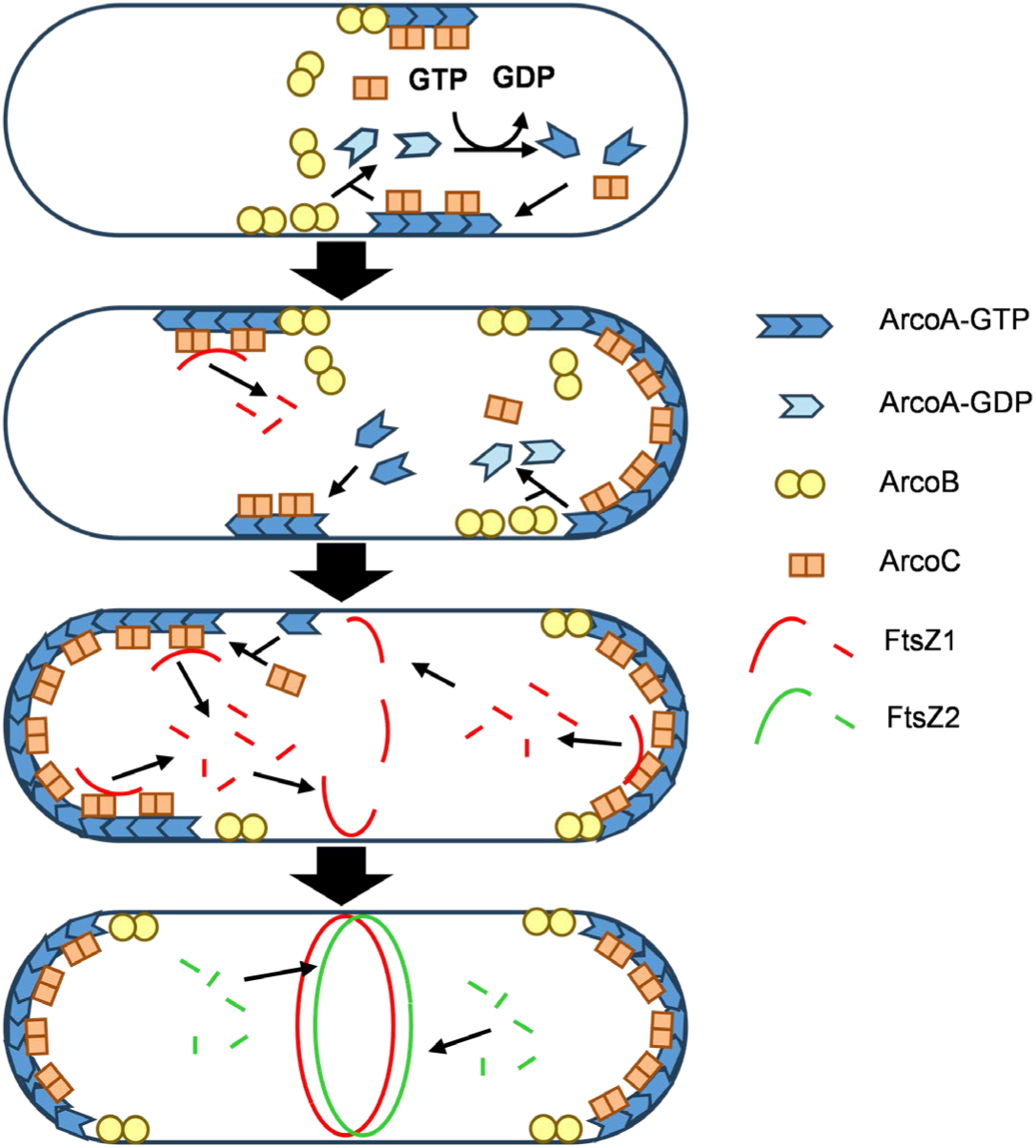
A working model for spatial regulation of cell division by the Arco oscillator in *H. volcanii*. ArcoA (an Era-like GTPase) binds to the membrane in its GTP-bound form via its N-terminal amphipathic helix. ArcoB (a SepF-like protein) also binds to the membrane via its amphipathic helix. ArcoA and ArcoB interact directly to generate the pole-to-pole oscillation. ArcoC (a zinc-finger protein with a long intrinsically disordered region) oscillates together with ArcoAB by interacting with ArcoA. At cell poles where the ArcoC concentration is high (due to the oscillation-generated concentration gradient), ArcoC binds directly to FtsZ1 and blocks its polymerization, preventing Z-ring assembly at the poles. At the cell center, where the ArcoC concentration is low, FtsZ1 is able to polymerize and recruits the other division proteins to assemble into the complete divisome, thereby establishing a correct division plane and driving cell constriction.

Intriguingly, the archaeal Arco system and the bacterial Min system share no sequence homology yet perform analogous functions: both generate a spatial gradient of an FtsZ inhibitor (ArcoC or MinC) that is high at poles but low at midcell, restricting Z-ring formation to the midcell. The bacterial Min system has diversified in different bacterial species. In *E. coli*, MinD and MinE oscillate pole-to-pole to generate the MinC gradient, whereas in *B. subtilis*, MinC and MinD are persistently anchored to poles by the curvature-sensing protein DivIVA and the adaptor MinJ, without an oscillation (Fig. 1a). Interestingly, global analysis of the Arco system across the archaeal domain showed that it has also diversified as the three genes are not always encoded in a single operon and some archaeal species contain only homologs of ArcoA and ArcoC. It would be interesting to explore what alternative factors are involved in regulating these variants containing only ArcoA and ArcoC, and whether they regulate cell division by forming bipolar gradients without apparent oscillations.

ArcoA, the primary member of the Arco system that likely hydrolyses GTP to provided energy for sustaining the oscillation, is an uncharacterized family of archaeal GTPases belong to the broader Ras superfamily GTPases. Ras and Ras-like proteins form various spatial patterns in eukaryotes, such as chromosomal gradients of Ran that guide microtubule nucleation ^39,40^, Cdc42 gradient that guides polarized growth in yeast ^41^, and Rac1 gradient for cell migration ^42^. Yet ArcoA contains a unique N-terminal amphipathic helix and is phylogenetically more closely related to the eukaryotic SRβ (transmembrane^43^) and bacterial MglA (cytosolic^44^), indicating that GTPases are capable of independently evolving diverse patterns. Archaea, especially Asgard archaea are known to encode a diverse array of GTPases with unknown functions ^45–47^. It would thus be intriguing to imagine a broader range of spatial patterns that may be fueled by these GTPases as well as other NTPases across the domain of archaea.

Notably, while we were preparing the manuscript, He and colleagues reported the discovery the Arco system independently^48^, which they named as the Dip (Divisome positioning) system. In agreement with our study, they found that the Dip proteins oscillate within the cells and regulate Z ring formation. They also demonstrated that DipA (ArcoA) is a membrane-binding GTPase with intrinsic GTPase activity, and DipB (ArcoB) can stimulate DipA’s GTPase activity and displaces it from the membrane ^48^. Thus, their biochemical data validate the GTPase cycle of AcroA and the antagonistic role of ArcoB underlying the oscillation. Overall, the two works are highly similar and complementary.

## Materials and Methods

### Strains and growth conditions

All strains used in this study are provided in Supplementary Table 6. *E. coli* strains JS238, JM110, and BL21(DE3) were used for plasmid propagation, generation of methylation-deficient DNA, and protein expression, respectively. These strains were grown in Luria-Bertani (LB) broth (10 g/L tryptone, 5 g/L yeast extract, 5 g/L NaCl, 0.05 mg/mL thymine) or on LB agar plates (1.5% agar) at the indicated temperatures. Ampicillin (100 μg/mL) was supplemented as needed to select for plasmid retention. *H. volcanii* strains were routinely cultured at 45 °C in Hv-YPCTE or Hv-Cab medium as described previously ^19^. When necessary, counter-selection was performed by adding 5-fluoroorotic acid (FOA) at 100 μg/mL. Expression from the *P_tna_* promoter was induced by adding L-tryptophan (Trp) at the indicated concentrations. Cell growth was followed by measuring the optical density at 600 nm (OD_600_). Unless otherwise noted, cultures were maintained in exponential phase (OD_600_ < 0.8) for at least two days prior to sample collection.

### Plasmid construction

All plasmids used in this study are listed in Supplementary Table 7, and the corresponding primers are provided in Supplementary Table 8. For plasmids used for genome editing, inducible expression, and split-GFP assays in *H. volcanii*, the backbone vectors pTA131^49^, pTA962^50^, and pTA1228^51^ were employed, respectively. For plasmids used for complementation experiments, each gene was cloned into the corresponding plasmids under its native promoter. Plasmids expressing GFP or mCherry fusions of FtsZ1, FtsZ2 and ArcoC were constructed by cloning the respective gene under the control of their respective native promoters. Plasmids expressing GFP fusion of ArcoB was constructed by cloning *arcoB* under the control of the xylose promoter (*P_xyl_*). For all other genes, GFP fusions were driven by the *P_tna_* promoter ^52^. To generate GFP or mCherry fusions, the open reading frame of the target gene was amplified and inserted in-frame with the fluorescent reporter gene. To build dual-expression plasmids, a DNA fragment carrying one gene was ligated into the NotI-linearized site of a plasmid that already contained the expression cassette for the other gene. For split-GFP assays, the sfGFP10 and sfGFP11 fragments were fused to the proteins of interest via flexible linkers (15–30 amino acids)^30^. Plasmids for protein purification were constructed using the pE-SUMO backbone. All DNA fragments were amplified by PCR using a high-fidelity DNA polymerase with the primers listed in Supplementary Table 8, purified by gel extraction, and then assembled into restriction enzyme linearized target vectors using a commercial homologous recombination cloning kit. All constructs were confirmed by Sanger sequencing, demethylated in *E. coli* JM110, and finally introduced into *H. volcanii* via PEG-mediated spheroplast transformation^49^.

### Genomic modifications

A two-step homologous recombination (pop-in/pop-out) strategy ^52^ was employed to generate gene deletion and promoter-replacement strains. To delete the *arcoC* gene in *H. volcanii*, a non-replicating plasmid carrying the upstream and downstream flanking regions of the *arcoC* start codon was first demethylated and then introduced into strain H26 via PEG-mediated spheroplast transformation. Transformants in which the plasmid integrated via a single-crossover event (pop-in) were selected on Hv-Cab agar plates lacking uracil. Subsequent excision of the plasmid (pop-out) was selected on Hv-YPCTE plates containing 5-fluoroorotic acid (FOA). The intended chromosomal modifications were confirmed by PCR followed by Sanger sequencing. The same procedure was used to generate deletion strains for *ddfA*, *ftsZ2*, *arcoA*, *arcoB*, and the *arco* operon in the H26 background. For the construction of *arcoA* L4* stain, a non-replicating plasmid carrying a mutant *arcoA* allele flanked by its upstream and downstream regions was introduced into an *arcoA* deletion strain, following the same procedure described above. The *arcoB* A4* strain was generated using a similar approach.

### Crosslinking and IP-MS to screen for division proteins and regulators

*H. volcanii* H26 expressing FtsZ1-GFP or FtsZ2-GFP under the control of the *P_tna_* promoter served as the experimental groups, while the same strain expressing only GFP was used as the negative control. Single colonies were picked into 5 mL of Hv-YPCTE medium and incubated at 45 °C for 48 h. The cultures were subsequently diluted 1:100 (v/v) into 80 mL of fresh Hv-YPCTE medium containing 1 mM Trp (three biological replicates per group) and grown at 45 °C for about 12 h until OD_600_ reached approximately 0.5. Cells were pelleted by centrifugation at 10,000 rpm for 10 min at 4 °C, then washed with 16 mL of high-salt PBS buffer (HSPB; PBS containing 144 g/L NaCl). For crosslinking, 640 μL of 25 mM DSP (3,3 ′ -dithiobis(succinimidyl propionate)) dissolved in DMSO was added to the cell pellet, followed by a 2 h incubation on ice. The crosslinking reaction was terminated by adding 2 mL of 1 M Tris-HCl (pH 8.2), gently mixed and kept at room temperature for 15 min. After a single wash with 16 mL of HSPB, the pellet was resuspended in 1 mL of HKPBT buffer (HKPB with 0.5% Tween-20; HKPB: PBS containing 194 g/L KCl) and disrupted by sonication. The lysate was clarified by centrifugation, and 200 μL of the cleared supernatant was set aside as the input sample for Western blotting to evaluate crosslinking efficiency. For immunoprecipitation, 800 μL of the cleared lysate was mixed with an anti-GFP antibody (1:200 dilution) and incubated at 4 °C for 8 h with gentle rotation. Meanwhile, 80 μL of magnetic beads were pre-washed twice with 800 μL of PBST (PBS with 0.5% Tween-20) and twice with HKPBT. The lysate-antibody mixture was then combined with the pre-washed beads and kept at 4 °C overnight. After incubation, the supernatant was removed, and the beads were washed three times with HKPBT and three times with HKPB (without Tween-20). One quarter of the beads were kept for efficiency testing, while the remaining three quarters were briefly dried by aspiration and stored at –80 °C for mass spectrometry analysis.

For Western blot analysis, the reserved beads (one quarter) were eluted with 30 μL of 1× SDS-PAGE loading buffer. A 10 μL portion was taken as the non-reduced sample. The remaining 20 μL was supplemented with 1 μL of 1 M DTT (final concentration ∼50 mM), heated at 95 °C for 10 min to reduce and reverse crosslinks, and used as the reduced sample. Input, post-IP supernatant, non-reduced eluate, and reduced eluate from each replicate were analyzed by immunoblotting to confirm successful immunoprecipitation and reversal of crosslinks. If the efficiency test showed no problems, the frozen beads were shipped to a commercial mass spectrometry facility (SpecAlly Life Technology Co. https://www.spec-ally.com/) for LC-MS/MS analysis.

Proteins exhibiting a fold change greater than 8 and a P-value less than 0.05 between the bait IP and the control were considered as candidate interactors of the bait protein. For each candidate interactor, two separate plasmids were constructed to produce N-terminal and C-terminal GFP fusions, respectively. The localization of each fusion was examined in H26 cells and proteins showing midcell localization were selected for further study.

### Microscopic observation

Prior to microscopy, all strains were subcultured twice overnight in Hv-Cab medium at 45 °C with shaking at 200 rpm. For strains in which gene expression was under the control of the *P_tna_* promoter, 0.2 mM Trp was added to induce the respective protein during the overnight incubation. Cells were collected at mid-exponential growth phase (OD_600_ ∼0.2-0.4). A 2 μL aliquot of the culture was placed onto a 1.5% agarose pad prepared with 18% BSW (Hv-Ca without casamino acids and CaCl_2_) and then covered with a glass coverslip. Fluorescence images were captured using an Olympus BX53 upright microscope fitted with a Retiga R1 camera, a CoolLED pE-4000 illumination source, and a U Plan XApochromat 100× oil-immersion objective (numerical aperture 1.45). GFP and mCherry signals were detected using Chroma filter sets 49002 and 49008, respectively. Image processing was carried out with Fiji, Adobe Photoshop 2021, or Adobe Illustrator 2023.

### Split-FP assays

Cells harboring split-GFP plasmids were subcultured twice overnight in Hv-Cab medium at 45 °C with shaking at 200 rpm to an OD_600_ of approximately 0.5. Expression was then induced by adding 0.4 mM Trp, and the cultures were shifted to 30 °C and incubated overnight with shaking at 200 rpm. Fluorescence signals were visualized by microscopy as described above. For quantitative analysis, the cell density of each culture was adjusted to an OD_600_ of 1.0, and 200 μL aliquots were transferred to a 96-well plate. Fluorescence intensity was measured using a Varioskan LUX plate reader (excitation at 489 nm, emission at 514 nm). For each experimental condition, two biological replicates and three technical replicates were performed. Statistical significance was assessed using a two-tailed Student’s *t*-test.

### 2D SIM fluorescence microscopy

For 2D structured illumination microscopy, strains were first transferred to Hv-Cab medium and grown twice overnight at 45 °C with shaking at 200 rpm. Expression was then induced by adding 1 mM Trp or 0.5 mM xylose, and the cultures were kept at 30 °C with shaking at 200 rpm overnight. Cells were collected during mid-exponential growth (OD_600_ ∼0.2-0.4). A 3.5 μL aliquot was placed onto a 1.5% agarose pad prepared with 18% BSW in a glass bottom dish (Cellvis). Imaging was performed using a HIS-SIM (High Intelligent and Sensitive Structured Illumination Microscopy) system (CSR Biotech) operating in 2D-SIM mode, with a 488 nm laser and a ×100/1.47 NA oil-immersion objective. Raw images were processed sequentially by Wiener and Sparse reconstruction to generate super-resolution images. The resulting super-resolution images were further processed using Fiji software.

### Time-lapse fluorescence microscopy

For time-lapse illumination microscopy, strains were subcultured twice overnight in Hv-Cab medium at 45 °C with shaking at 200 rpm. 1 mM Trp or 0.5 mM xylose was added during the overnight incubation for induction. Cells were collected during mid-exponential growth (OD600 ∼0.2-0.4). A 3.5 μL aliquot was placed onto a 1.5% agarose pad prepared with 18% BSW in a glass bottom dish (Cellvis). Imaging was performed using a HIS-SIM system (CSR Biotech) operating in widefield mode, with a ×100/1.47 NA oil-immersion objective. For time-lapse imaging of GFP or GFP fusions, cells were illuminated with a 488 nm laser, and images were captured every 20 seconds for a total duration of 10 minutes. For dual-color imaging of GFP and mCherry fusions, both 488 nm and 561 nm lasers were used, and images were acquired every 40 seconds over a 10-minute period. Time-lapse images were subsequently processed using Fiji software.

### Western blotting

Protein samples were resolved by SDS-PAGE on 12% polyacrylamide gels and then electrotransferred onto nitrocellulose (NC) membranes (Pall). After blocking with 5% (w/v) non-fat milk for 1 h at room temperature, the membranes were incubated overnight at 4 °C with the respective primary antibodies (diluted 1:10,000). Following five washes with TBST, the blots were treated with HRP-conjugated secondary antibodies (diluted 1:10,000) for 1 h at room temperature. Chemiluminescent signals were developed using an ultra-sensitive ECL substrate and captured with a ChemiDoc imaging system.

### Co-immunoprecipitation

Cells expressing the appropriate tagged proteins were cultivated in 40 mL of Hv-YPCTE medium (or Hv-cab medium when *P_xyl_* was uesd) at 45 °C with orbital shaking at 200 rpm until the culture reached an OD_600_ of about 1.0. The cultures were then switched to 30 °C, shaking at 200 rpm, and induced overnight by the addition of 1 mM Trp or 0.5 mM xylose. After induction, cells were collected by centrifugation at 10,000 rpm for 10 min at 4 °C. The resulting pellet was suspended in 2 mL of HKPB buffer containing a protease inhibitor cocktail and disrupted by sonication. The lysate was clarified by centrifugation under the same conditions 10,000 rpm for 10 min at 4 °C. For the input fraction, 200 μL of the cleared supernatant was combined with 50 μL of 5× SDS-PAGE loading buffer and heated at 95 °C for 10 min. In parallel, 400 μL of the supernatant was incubated overnight at 4 °C with magnetic beads pre-coated with anti-GFP, anti-Flag, anti-HA or anti-His antibodies (as specified). The beads were then washed three times with HKPBT, and bound immune complexes were eluted by boiling in 1× SDS-PAGE loading buffer. Both the eluted samples and the input samples were subsequently analyzed by Western blotting as described in the previous section.

### Expression level analysis

*H. volcanii* H26 strains harboring plasmids that encode GFP fusions of either the wild type or truncated versions of the target proteins, all under the transcriptional control of *P_tna_* or *P_xyl_* promoter, were inoculated into 40 mL of Hv-YPCTE medium. The cultures were incubated at 45 °C with shaking at 200 rpm for 16 h. Following this, gene expression was induced by supplementing the medium with 1 mM Trp or 0.5 mM xylose, and the cultures were shifted to 30 °C and allowed to grow overnight with continued shaking at 200 rpm. After determining the OD_600_ of each culture, the cell densities were adjusted to uniformity across all samples. An equal volume of each culture was then collected by centrifugation at 10,000 rpm for 10 min at 4 °C. The resulting pellets were washed once with 10 mL of HSPB and subsequently resuspended in 2 mL of HKPB containing a protease inhibitor cocktail. The cells were disrupted by sonication, and the lysates were clarified via centrifugation. For sample preparation, 200 μL of the cleared supernatant was mixed with 50 μL of 5× SDS-PAGE loading buffer and heated at 95 °C for 10 min. The prepared samples were then subjected to Western blot analysis as previously described.

### Protein expression and purification

Recombinant proteins were produced in *E. coli* BL21(DE3) transformed with plasmids pZA111 (H-*Af*FtsZ1), pZA113 (H-*Af*ArcoA), pZA116 (H-SUMO-*Af*ArcoB), or pZA106 (H-SUMO-*Af*ArcoC). Transformed cells were first grown overnight in LB medium containing 100 μg/ml ampicillin. The next day, each culture was diluted 1:100 into 300 ml of fresh LB and incubated at 37 °C until the OD_600_ reached 0.4. Protein expression was then induced by the addition of IPTG to a final concentration of 1 mM, and the cultures were allowed to grow for an additional 3 h at 37 °C. Cells were harvested by centrifugation and stored at −80 °C.

For purification, frozen cell pellets were thawed, resuspended in 20 ml of lysis buffer (50 mM Tris-HCl [pH 8.0], 200 mM NaCl, 5% glycerol, 10 mM imidazole), and lysed by sonication. The lysates were centrifuged at 18,000 g for 10 min at 4 °C to remove insoluble debris. The cleared supernatants were then loaded onto a pre-equilibrated Ni-NTA agarose column. After washing with wash buffer (50 mM Tris-HCl [pH 8.0], 200 mM NaCl, 5% glycerol, 20 mM imidazole), bound proteins were eluted with elution buffer (50 mM Tris-HCl [pH 8.0], 200 mM NaCl, 5% glycerol, 300 mM imidazole). Fractions containing the target proteins, as judged by SDS-PAGE, were pooled, dialyzed against storage buffer (50 mM Tris-HCl [pH 8.0], 200 mM NaCl, 5% glycerol), aliquoted, and frozen at −80 °C.

To remove the H-SUMO tag from *Af*ArcoB or *Af*ArcoC, the purified proteins were incubated with purified 6×His-tagged SUMO protease (Ulp1) at 30 °C for 1 h. The digestion mixture was subsequently passed through a fresh Ni-NTA column, which retained the released His-SUMO tag and the His-tagged protease, while the untagged target proteins were collected in the flow-through. After dialysis against storage buffer and concentration, the tag-free proteins were stored at −80 °C.

### Biolayer interferometry assays

To evaluate three pairwise interactions, namely *Af*ArcoC with *Af*FtsZ1, *Af*ArcoA with *Af*ArcoC, and *Af*ArcoA with *Af*ArcoB, independent binding assays were conducted at 30 °C using an Octet-Red96 system. For each assay, Ni-NTA biosensors were first pre-equilibrated in 1×Pol buffer (12.5 mM HEPES-NaOH [pH 6.8], 12.5 mM KCl, 2.5 mM MgCl_2_, 0.02% Tween-20). The 6×His-tagged bait protein (0.5 μM) was subsequently applied to the sensor tips for 3 min in 200 μL of the same buffer supplemented with 1 mM GTP, followed by a 60-second wash to eliminate non-specifically bound proteins. The untagged prey protein was allowed to associate for 4 min under shaking at 1,000 rpm, and dissociation was monitored for 3 min in buffer without any protein. Specifically, the following bait-prey pairs were tested: His-*Af*FtsZ1 with untagged *Af*ArcoC, His-*Af*ArcoA with untagged *Af*ArcoC, and His-*Af*ArcoA with untagged *Af*ArcoB. Each pair was assayed for at least three times. Data analysis and curve fitting were carried out using the ForteBio Data Analysis Software and GraphPad Prism 10.

### Transmission electron microscopy

To examine whether *Af*ArcoC influences the polymerization of *Af*FtsZ1, we performed polymerization assays followed by negative-staining transmission electron microscopy. All reactions (50 µL final volume) were conducted at room temperature in Pol buffer (50 mM HEPES-OH (pH 6.8), 50 mM KCl, 10 mM MgCl_2_). Four reaction conditions were set up: (i) 2.5 μM His-*Af*FtsZ1, 5 μM untagged *Af*ArcoC, and 1 mM GTP; (ii) 2.5 μM His-*Af*FtsZ1 and 1 mM GTP; (iii) 2.5 μM His-*Af*FtsZ1 and 1 mM GDP; (iv) 5 μM untagged *Af*ArcoC and 1 mM GTP. After adding the nucleotide (GTP or GDP), the mixtures were incubated for 5 min. Subsequently, 15 μL aliquots were deposited onto glow-discharged carbon-coated copper grids. Following a 5 min adsorption period, excess liquid was removed by blotting filter paper, and the grids were stained with 15 μL of 1% (w/v) uranyl acetate for 1 min, followed by another blotting step. The grids were air-dried overnight (>12 h) and imaged using a JEM-1400Plus transmission electron microscope operating at 100 kV.

### Protein identification and dataset collection

To identify homologues of the ArcoABC system across archaea, we searched a protein dataset comprising 6,717 archaeal genomes from the Genome Taxonomy Database (GTDB; release 226)^53^, supplemented with 916 Asgard archaeal genomes^54^, resulting in a curated, nonredundant set of 7,633 genomes for taxonomic distribution analyses. Profile hidden Markov model searches were performed using HMMER v3.3.2 (http://hmmer.org/) with pfam profiles^55^ corresponding to ArcoA (PF01926), ArcoB (PF09846), and ArcoC (PF09845). Hits were retained when both the full-sequence and best-domain scores reached the gathering thresholds encoded in the respective profiles: 21.9 for ArcoA, 27.0 for ArcoB, and 23.0 for ArcoC. Protein identifiers were reconciled with their corresponding genome and gene-coordinate records by normalizing accession versions and zero-padded contig and gene indices.

To examine the distributions of FtsZ1, FtsZ2, and SepF across archaea, we retrieved the reference protein sequence datasets provided by Pende et al^13^. Separate profile hidden Markov models were constructed from the FtsZ1 and FtsZ2 reference sequence sets and used to search the same combined protein database. For visualization, each protein was classified as present in a lineage when detected in more than 50% of its genomes, partially present when detected in 10-50% of its genomes, and absent when detected in fewer than 10% of its genomes. The distributions were visualized together with the ArcoABC patterns and the archaeal reference phylogeny using iTOL.

### Gene-cluster and taxonomic distribution analyses

For each genome, retained ArcoA, ArcoB, and ArcoC homologues were mapped to their contigs and ordered by gene index to examine local gene organization. Components located on the same contig with no more than one intervening gene were considered directly linked, whereas those separated by two to five intervening genes were classified as gapped but linked. Components separated by more than five genes, located on different contigs, or lacking sufficient positional information were treated as dispersed. Genomes were accordingly classified as encoding a complete linked

ArcoABC cluster, a linked ArcoAB pair with a separated ArcoC homologue, a partial linked pair, dispersed components, a single detected component, or no detectable component. Taxonomic assignments were used to calculate the proportions of these genomic architectures within well-sampled archaeal lineages. The distributions of FtsZ1, FtsZ2, and SepF homologues were mapped onto the same lineage framework for comparison with that of ArcoABC.

### ArcoA phylogenetic analysis

To investigate the evolutionary relationship between ArcoA and other GTPase families, we compiled a dataset of 281 representative protein sequences. The 996 GTPase sequences underlying the phylogenetic analyses of Vargova et al.^45^ were retrieved from the associated public dataset and pruned to 250 representatives using Treemmer v0.3, with the target number of retained leaves set to 250 and all other parameters left at their defaults. ArcoA homologues encoded in ArcoABC gene clusters were retrieved from searches against the combined GTDB archaeal and supplemental Asgard protein database and clustered using MMseqs2^56^ v13.45111 with the parameters --min-seq-id 0.6 -c 0.8 --cov-mode 1. A preliminary phylogeny was constructed using FastTree^57^ v2.2.0 with default parameters, and the dataset was further reduced to 30 phylogenetically representative ArcoA sequences using Treemmer^58^ v0.3, with the target number of retained leaves set to 30 and all other parameters left at their defaults. The ArcoA protein from *Haloferax volcanii* was added separately. The resulting 281 sequences were aligned using MAFFT v7.526^59^ with the --auto option, and poorly aligned regions were removed using trimAl v1.4 rev build with the -gappyout option. Maximum-likelihood analysis was performed using IQ-TREE^60^ v3.0.1 under the LG+C60+F+G4 model with 1,000 ultrafast bootstrap^61^ replicates. The resulting tree was midpoint rooted and visualized using iTOL.

### ArcoB phylogenetic analysis

Based on the above sequence search, forty representative ArcoB homologues encoded in ArcoABC gene clusters and 290 sequences of SepF were respectively selected via MMseqs then added to ensure adequate representation of the operon-associated ArcoB family. The resulting dataset comprised 330 protein sequences. These sequences were aligned using MAFFT v7.526 with the --auto option, and poorly aligned regions were removed using trimAl v1.4 rev build with the -gappyout option. Maximum-likelihood analysis was performed using IQ-TREE v3.0.1 under the LG+C60+F+G4 model with 1,000 ultrafast bootstrap replicates. The resulting tree was midpoint rooted and visualized using iTOL.

### Statistical analysis

All statistical evaluations were carried out using GraphPad Prism 10. Comparisons between two groups were assessed with a two-tailed, unpaired Student’s *t*-test. A P-value below 0.05 was regarded as statistically significant. Each experiment was conducted at least three times independently, yielding comparable results. Quantitative data are expressed as mean ± standard deviation (SD). The exact sample sizes (n) for each experiment are provided in the corresponding figure legends.

### Reagents and chemicals

All reagents and chemicals are listed in Supplementary Table 9.

### Data availability

Data generated and analyzed during this study are presented in the paper or in the Supplemental Information and Datasets. Protein sequences, multiple alignment, and phylogenetic trees are provided in the Supplemental Information. Plasmids and strains that support the findings of this study are available from the corresponding authors on reasonable request.

## Acknowledgements

We thank members of the Du lab, Wu lab and Chen lab for advice and helpful discussions to carry out this study. This study was supported by National Natural Science Foundation of China (grants 32270049 and U24A20345), and the Young Top-notch Talent Cultivation Program of China to S.D.; National Natural Science Foundation of China (grant 323B2001) to S.Z.; Research in the Wu lab was supported by National Natural Science Foundation of China (grant 32570019 and 32370003).

## Supplementary Information include

### Supplementary Figures 1-9

**Fig S1.**
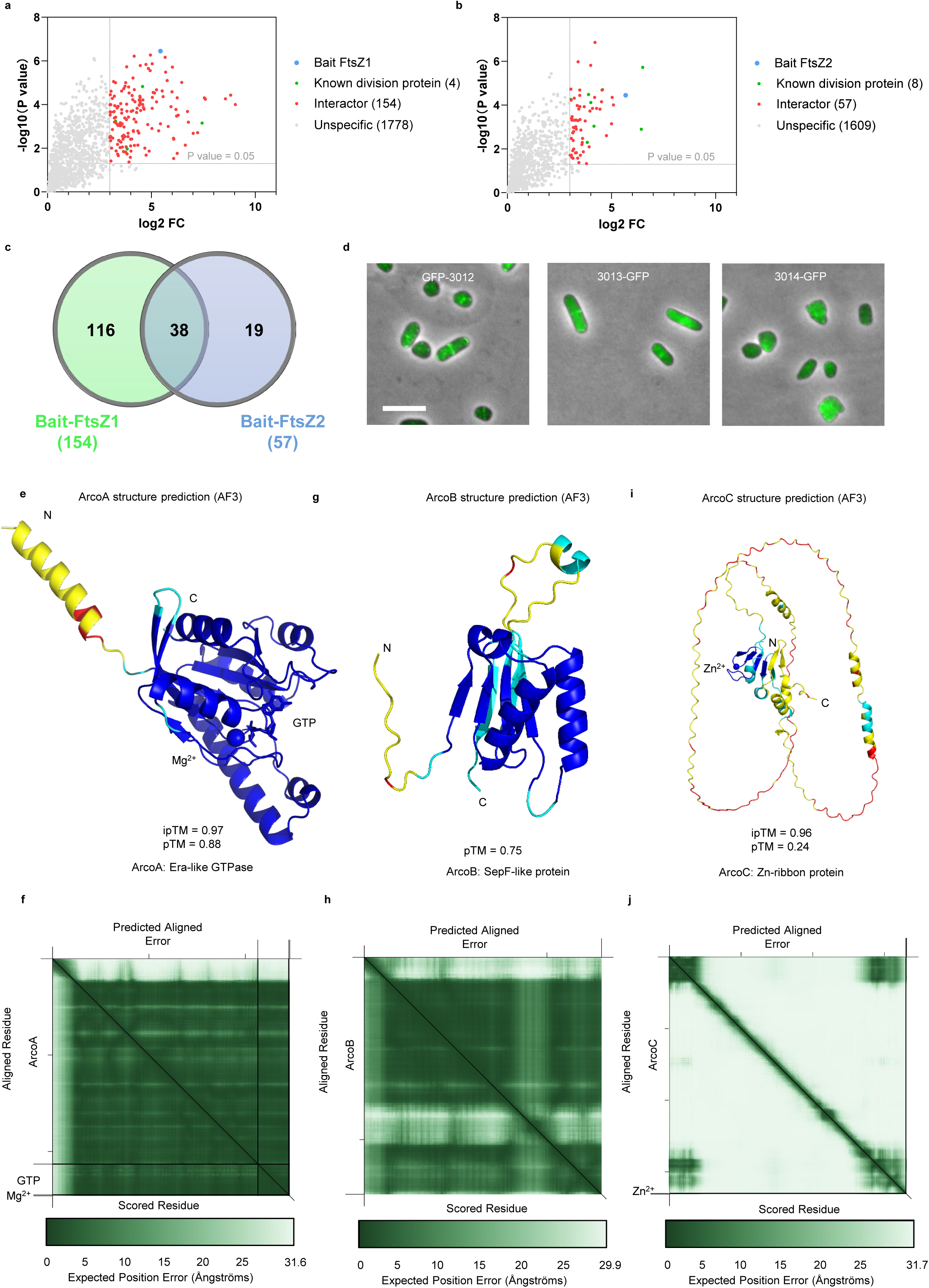
Identification of the Arco system in *H. volcanii*.

**Fig S2.**
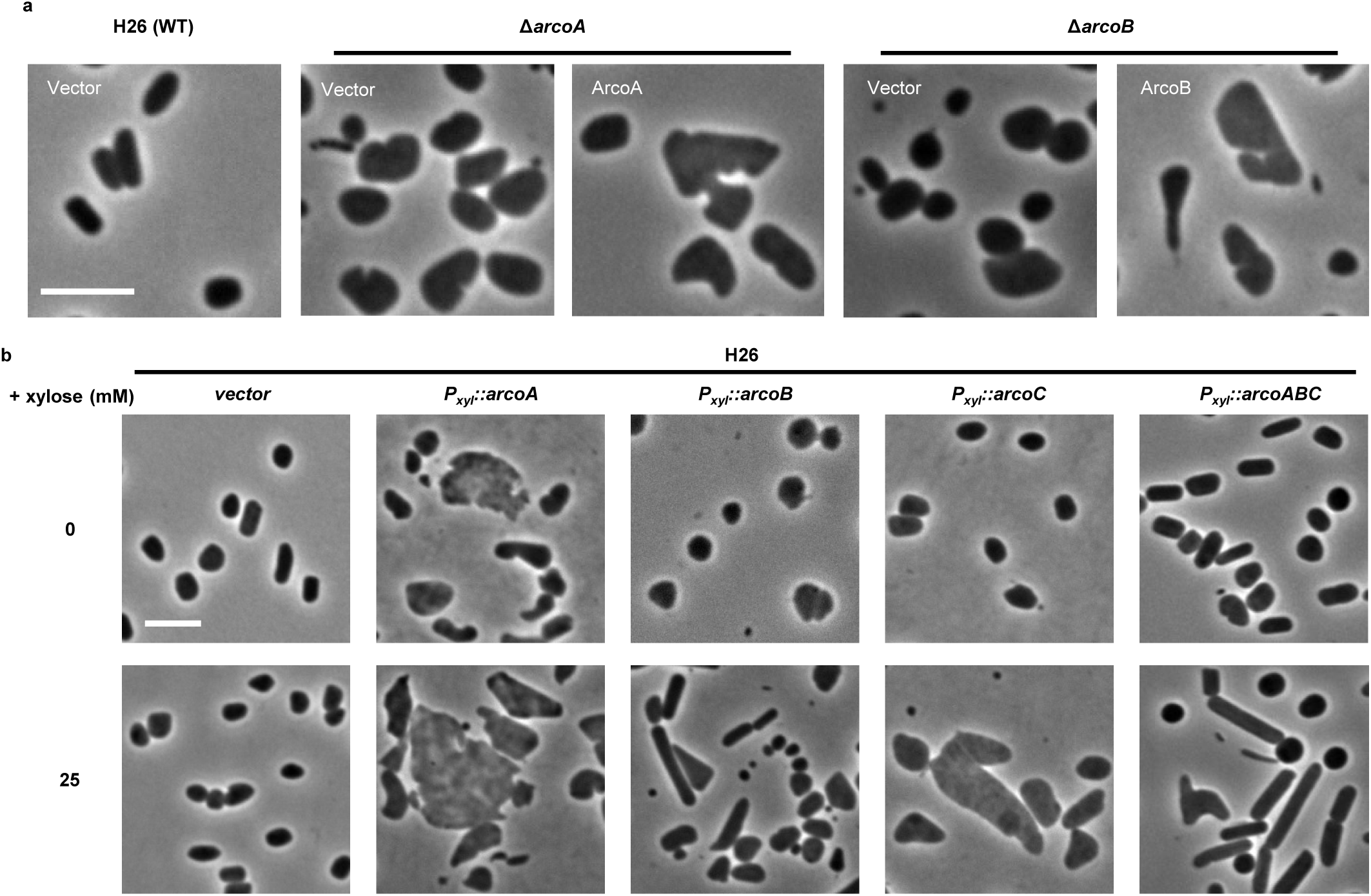
Phenotypes of the *arcoA* or *arcoB* deletion strain and overexpression of Arco proteins.

**Fig S3.**
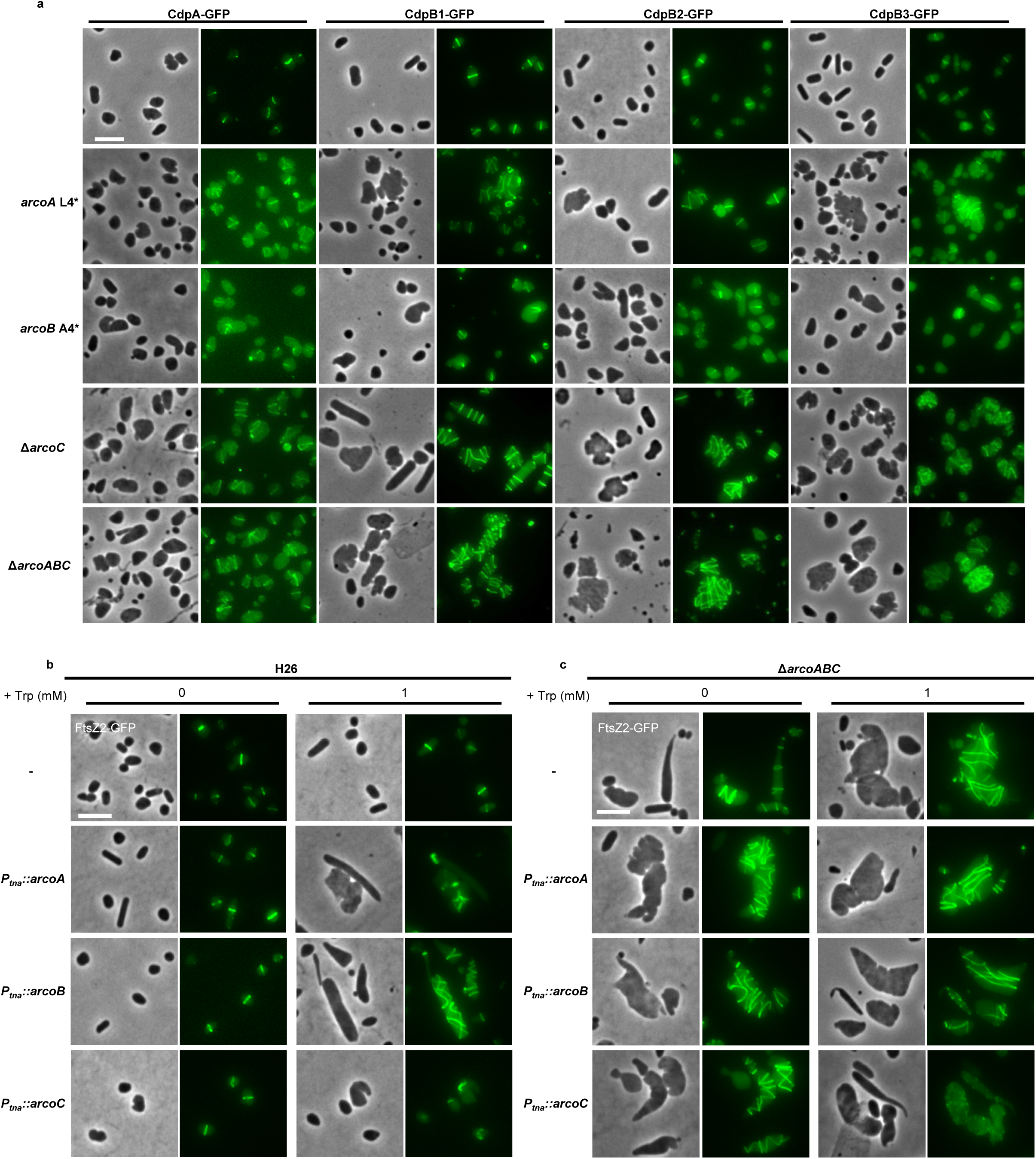
The Arco system controls localization of division proteins.

**Fig. S4.**
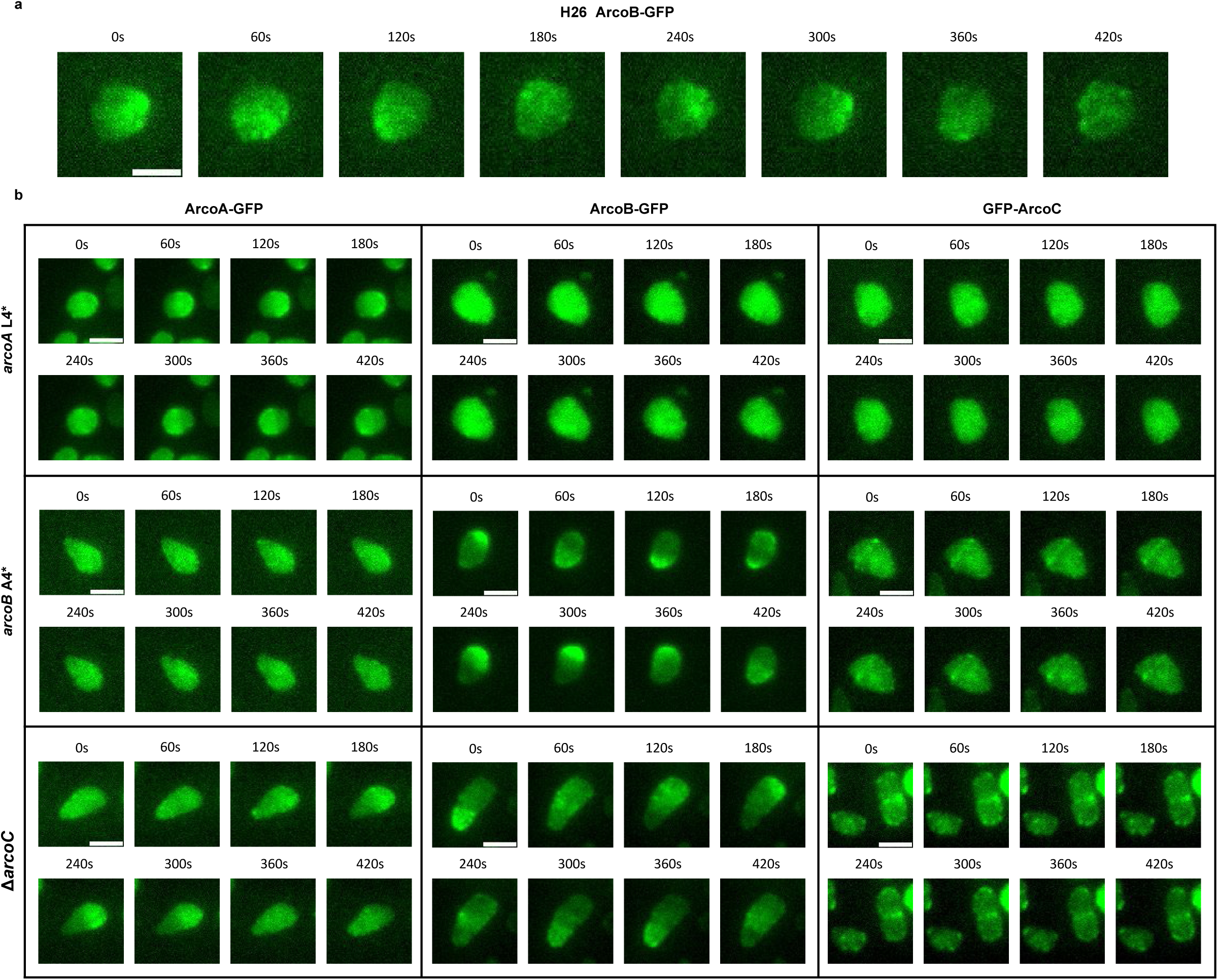
Oscillation modes and dependency of the Arco system.

**Fig. S5.**
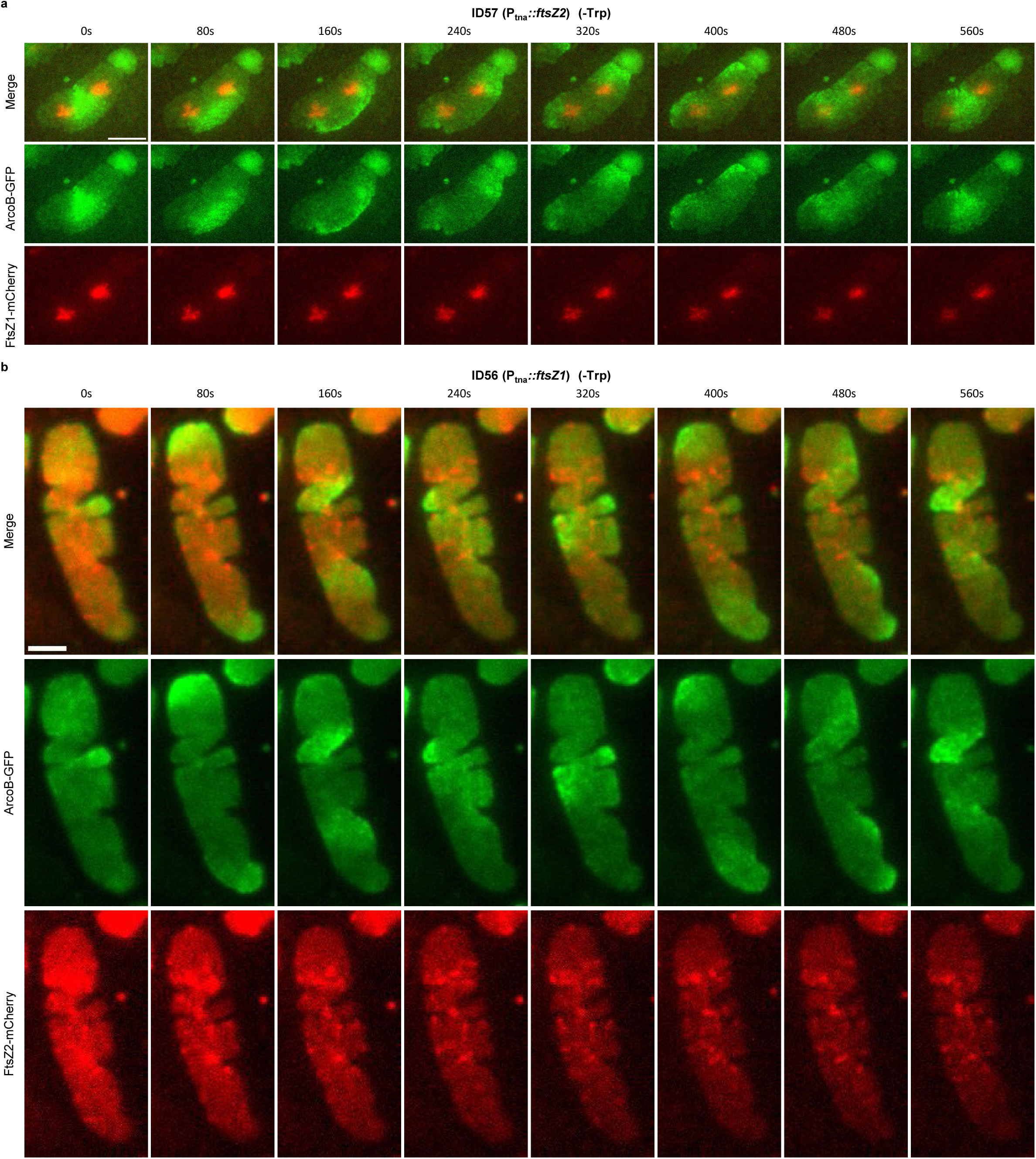
Arco oscillations restricts FtsZ1 localization but not FtsZ2.

**Fig. S6.**
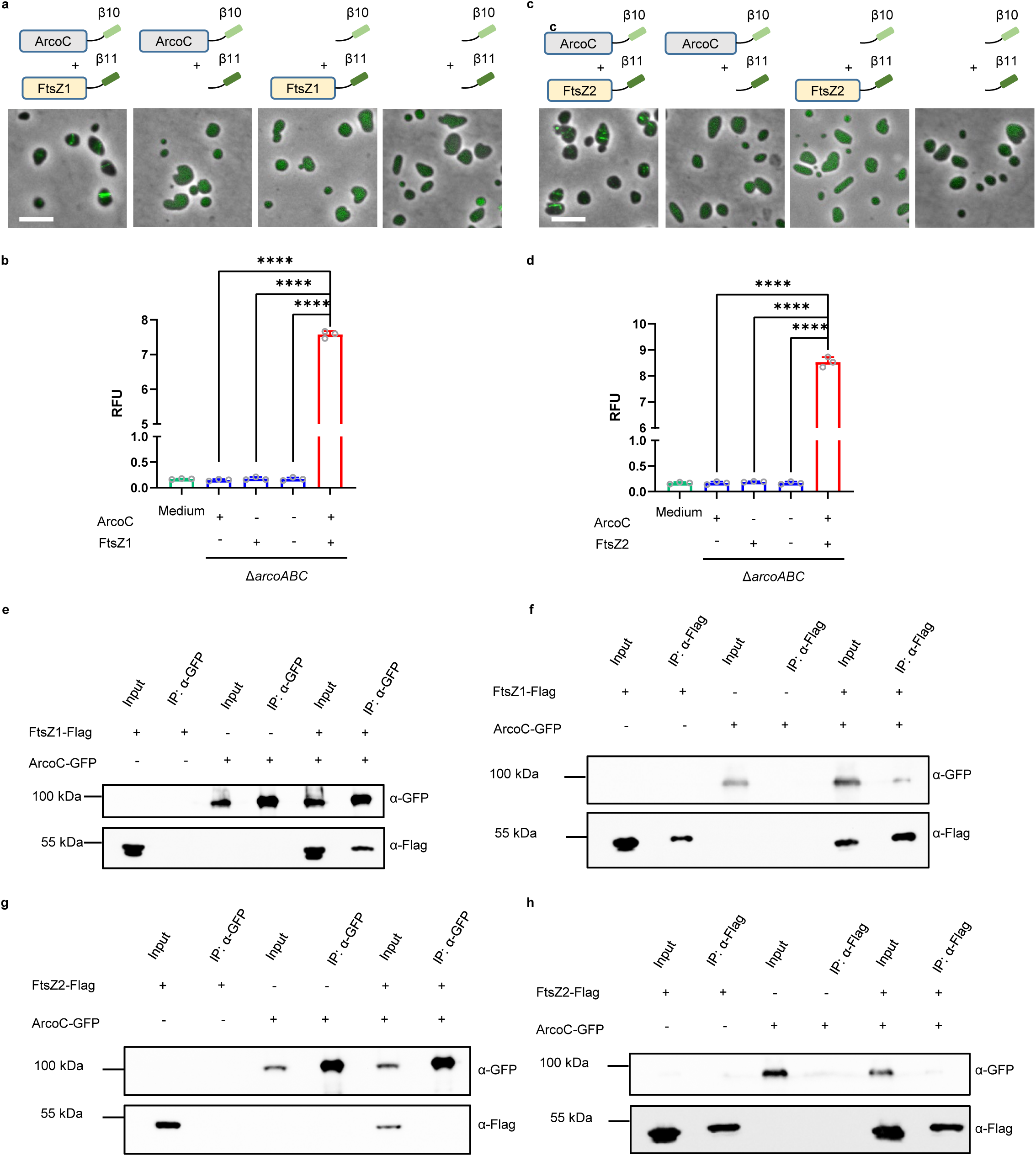
ArcoC interacts with FtsZ1 *in vivo*.

**Fig. S7.**
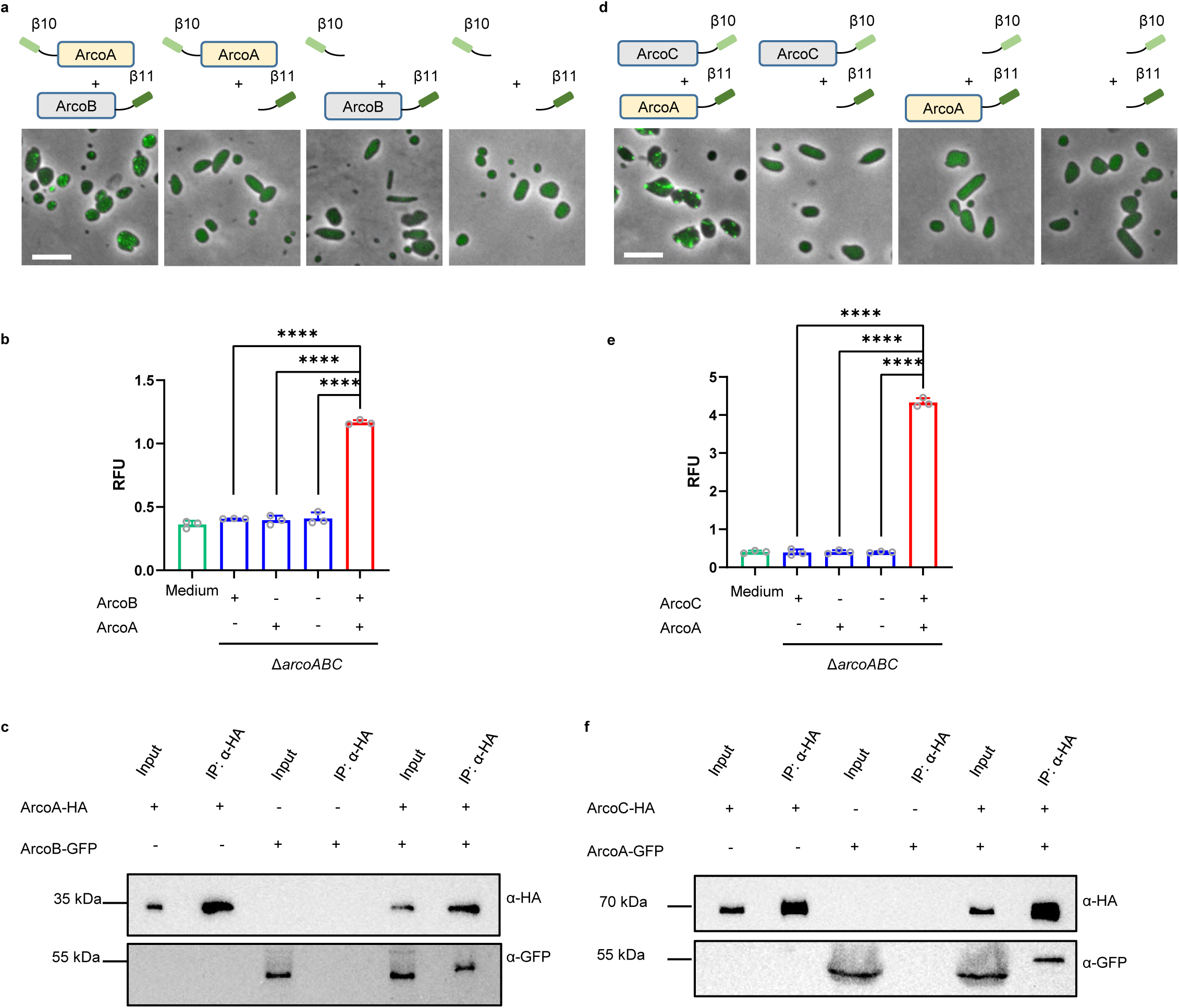
ArcoA interacts with both ArcoB and ArcoC *in vivo*.

**Fig. S8.**
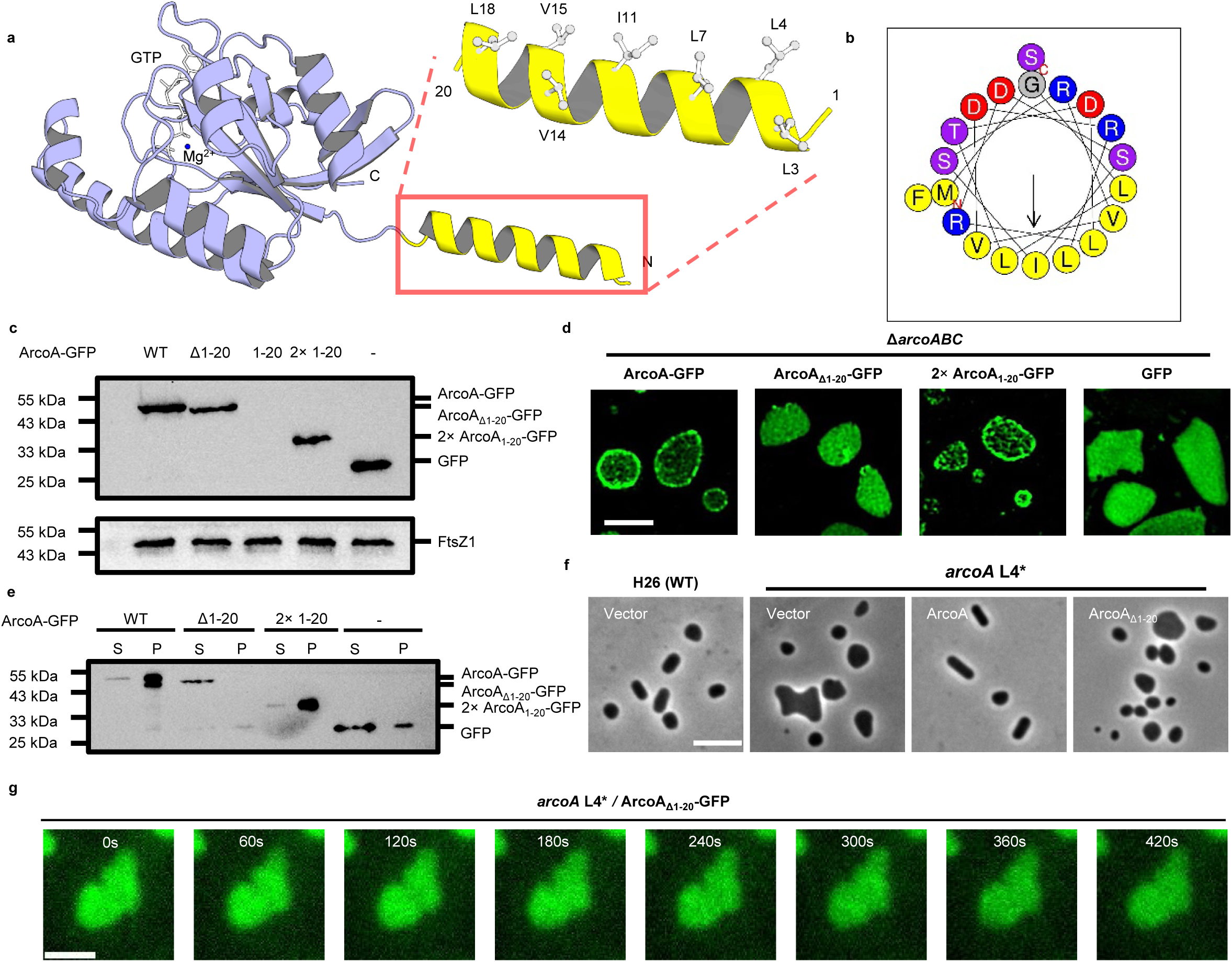
The amphipathic helix of ArcoA is essential for membrane binding and oscillation.

**Fig. S9.**
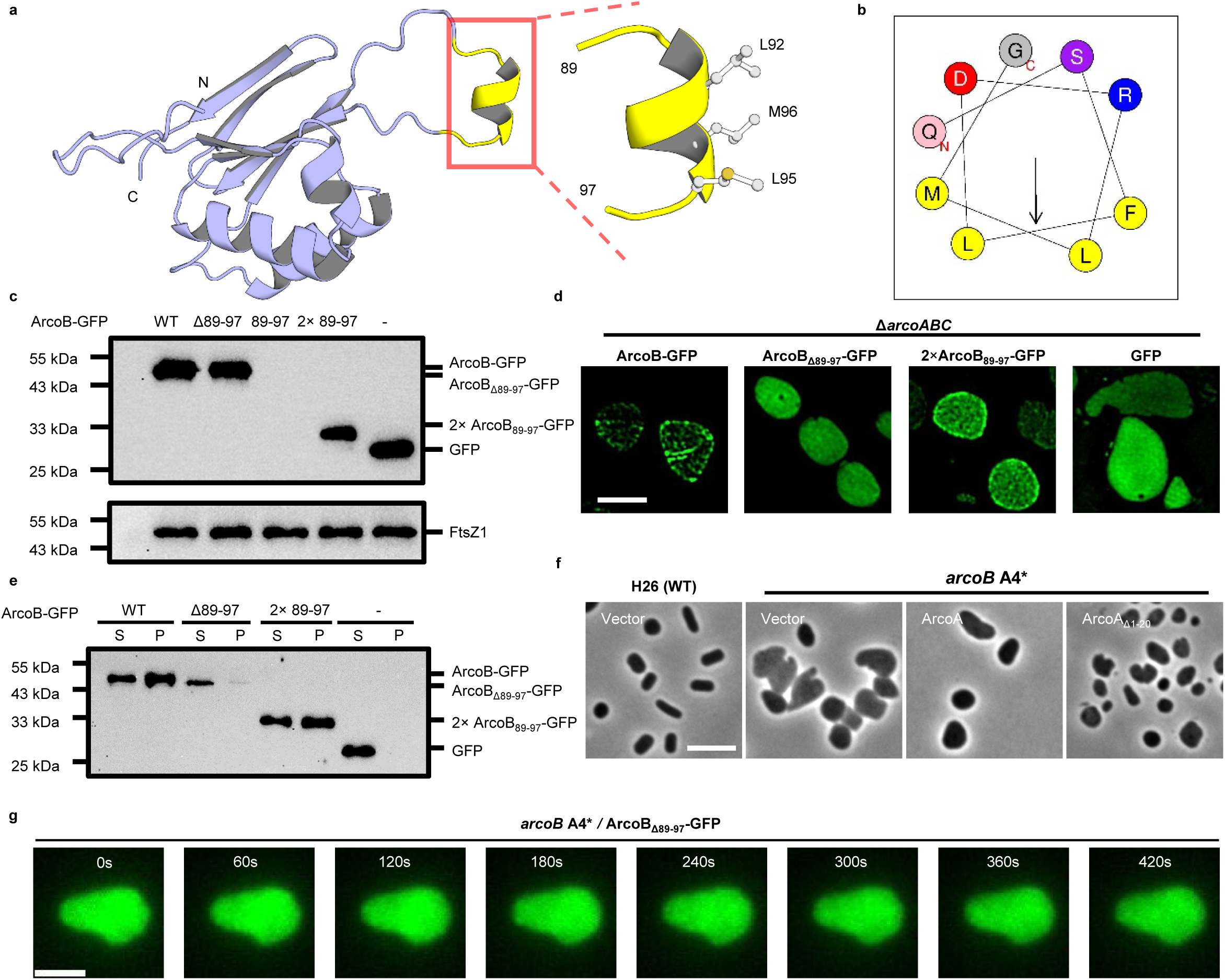
The amphipathic helix of ArcoB is essential for membrane binding and oscillation.

### Supplementary Table 1-9

Table S1. Proteins identified by in vivo crosslinking IP-MS targeting FtsZ1. Table S2. Proteins identified by in vivo crosslinking IP-MS targeting FtsZ2. Table S3. Identified potential interactors of both FtsZ1 and FtsZ2.

Table S4. Localization of the 38 common interactors of both FtsZ1 and FtsZ2. Table S5. Information of the Arco proteins.

Table S6. Strains used in this study. Table S7. Plasmids used in this study. Table S8. Primers used in this study.

Table S9. Reagents and chemicals used in this study. Table S10. Protein sequences and alignments.

### Supplemental Videos 1-10

Supplementary Video 1. Oscillation of ArcoA-GFP in rod-shaped wild type cells. Supplementary Video 2. Oscillation of ArcoB-GFP in rod-shaped wild type cells.

Supplementary Video 3. Oscillation of ArcoB-GFP in disc-shaped wild type cells. Supplementary Video 4. Oscillation of ArcoC-GFP in wild type cells.

Supplementary Video 5. Localization dynamics of ArcoB-GFP and FtsZ1-mCherry in rod-shaped wild type cells.

Supplementary Video 6. Localization dynamics of ArcoB-GFP and FtsZ1-mCherry in triangular wild type cells.

Supplementary Video 7. Localization dynamics of ArcoB-GFP and FtsZ1-mCherry in elongated and rod-shaped *ΔddfAΔftsZ2* cells.

Supplementary Video 8. Localization dynamics of ArcoB-GFP and FtsZ1-mCherry in FtsZ2 depleted cells.

Supplementary Video 9. Localization dynamics of ArcoB-GFP and FtsZ2-mCherry in FtsZ1 depleted cells.

Supplementary Video 10. Localization dynamics of ArcoA_Δ1-20_-GFP in ArcoA null (AcroA L4*) cells.

Supplementary Video 11. Localization dynamics of ArcoB_Δ89-97_-GFP in ArcoB null (AcroB A4*) cells.

