## Supplementary material for "A self-organized geometry sensing oscillator spatially regulates cell division in archaea": Fig. S1-9

**Affiliation**

### Supplementary Figure Legends

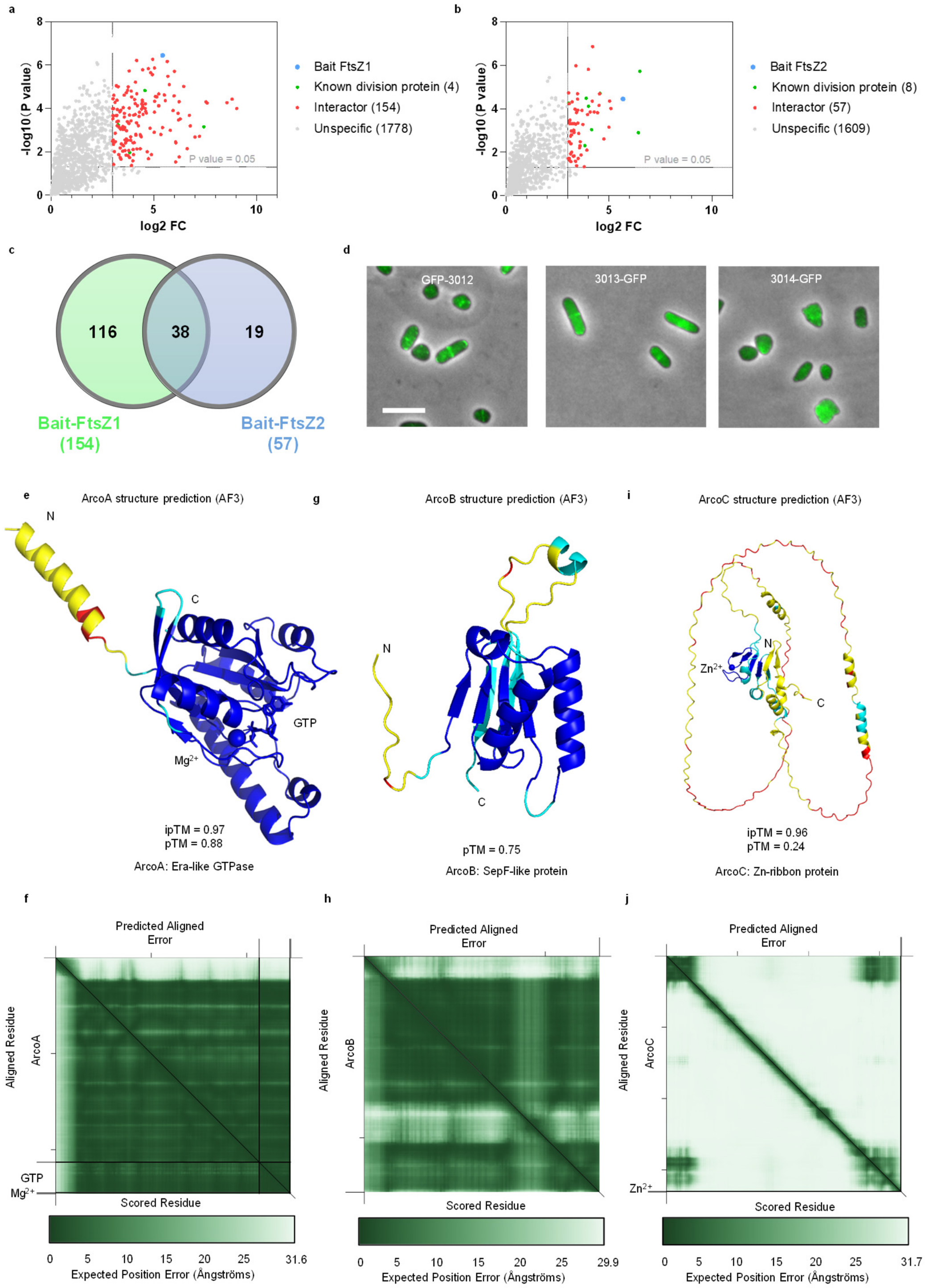


**Fig S1 Identification of the Arco system in *H. volcanii*.**

a-b. Volcano plot displays known division proteins and candidate interactors of FtsZ1 (a) or FtsZ2 (b) by CLIP-MS. Proteins with an enrichment ≥8-fold over the control and a P-value < 0.05 were considered as candidate interactors. 4 known division proteins (green) along with 154 additional interactors (red) were detected in the FtsZ1-GFP immunoprecipitate, whereas 8 known division proteins and 57 additional interactors were found in the FtsZ2-GFP sample.

c. Venn diagram showing overlap of FtsZ1 and FtsZ2 interactors. 38 common interactors are listed in Supplementary Table 3.

d. Representative images of localization of GFP-tagged ArcoA, ArcoB and ArcoC in wild type H26 cells. All GFP fusion proteins were expressed under the control of the $P_{tna}$ promoter from plasmids, induced overnight with 0.2 mM tryptophan (Trp) at 45 °C. Scale bar, 5 μm.

e-f. Structural model of ArcoA with GTP and magnesium ion colored by model confidence (blue - pLDDT > 90%, cyan - 90% > pLDDT > 70%, yellow – 70% > pLDDT > 50%, red - pLDDT <50%), ipTM = 0.97, pTM=0.88. Predicted aligned error (%) heatmap plot (f). The degree of confidence in pairwise interactions between residues was shown.

g-h. Structural model of ArcoB colored by model confidence (g) and predicted aligned error (%) heatmap plot (h). Parameters were similar as in e-f.

i-j. Structural model of ArcoC with one bound zinc ion colored by model confidence (i) and predicted aligned error (%) heatmap plot (i). Parameters were similar as in e-f.

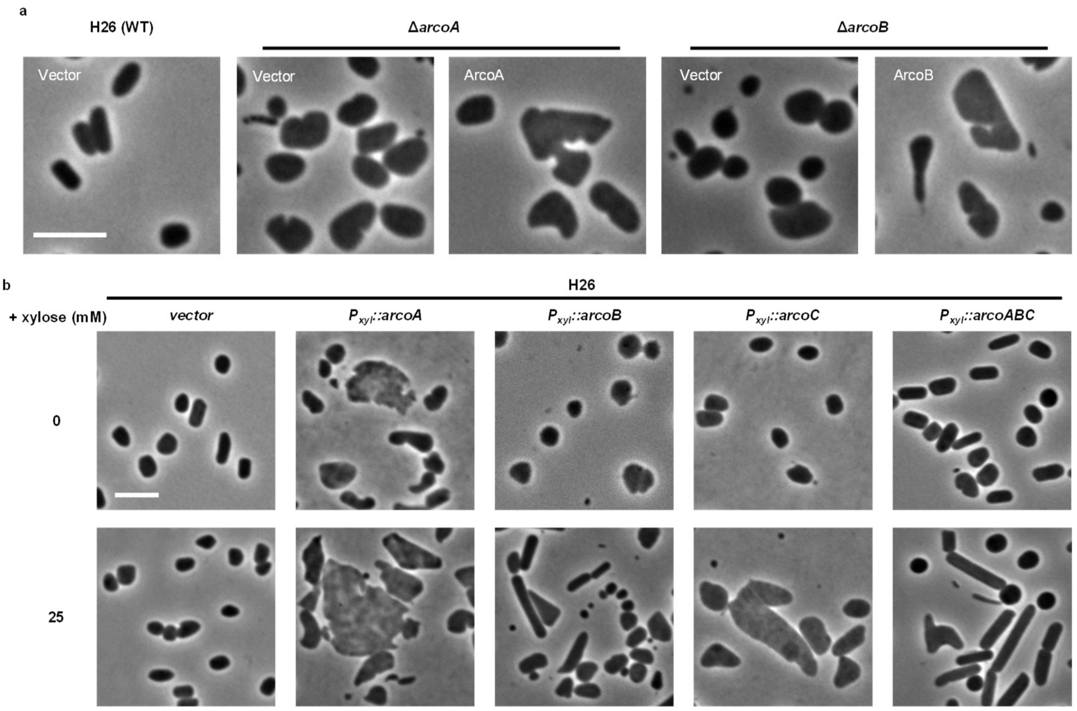


**Fig S2. Phenotypes of the *arcoA* or *arcoB* deletion strain and overexpression of Arco proteins.**

a. Representative phase-contrast images of wild type H26, Δ*arcoA*, and Δ*arcoB* cells with or without plasmid-borne complementation. The corresponding gene was expressed from its native promoter on a plasmid. Scale bar, 5 μm.

b. Representative images of wild type cells overexpressing individual Arco proteins or the full operon. ArcoA, ArcoB, ArcoC or the full operon were expressed from a plasmid driven by $P_{xyl}$ promoter, induced with indicated concentration of xylose. All strains were grown at 45 °C overnight before imaging. Scale bar, 5 μm.

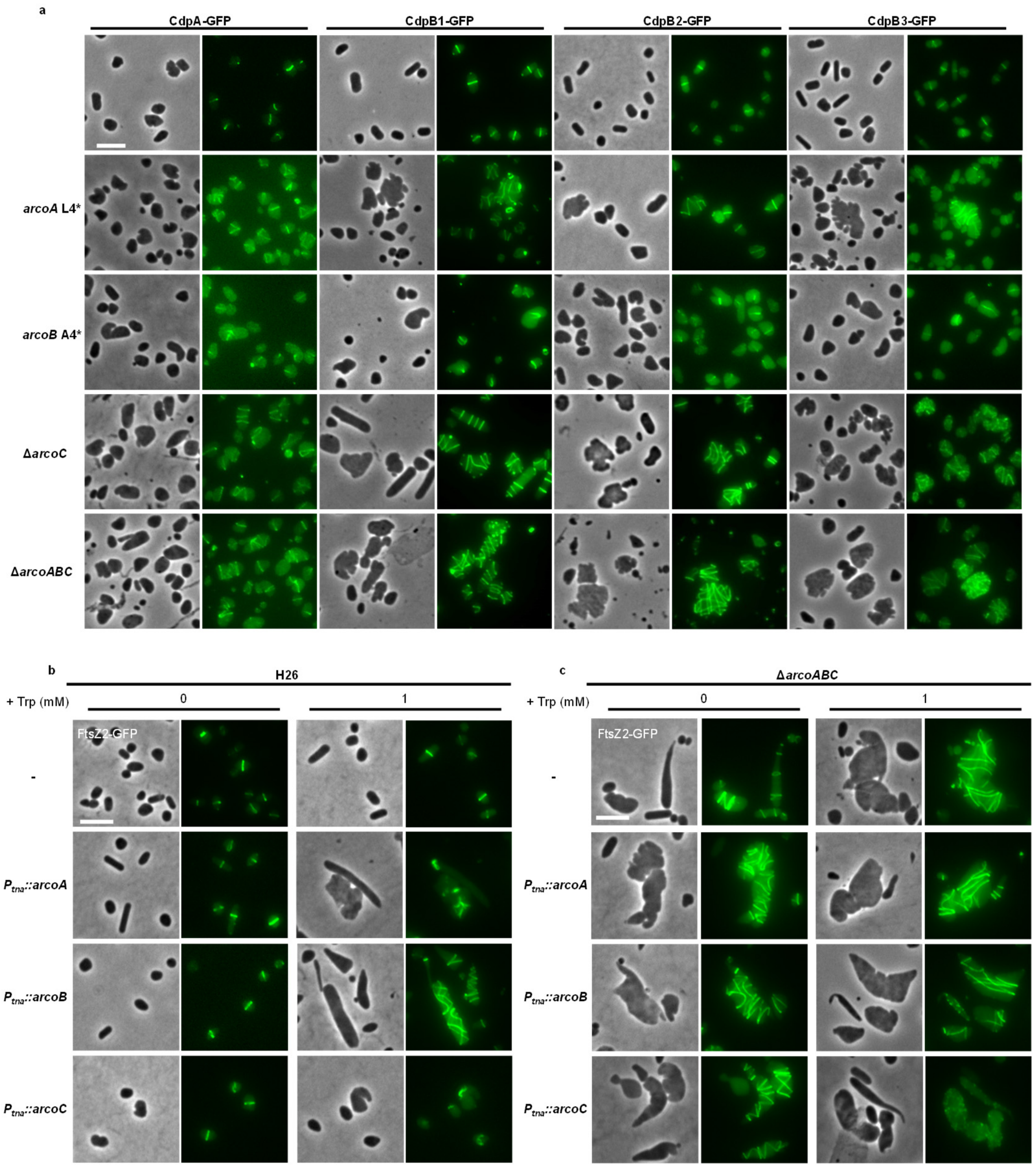


**Fig S3 The Arco system controls localization of division proteins.**

a. Representative images of the localization of GFP fusions of CdpA, CdpB1, CdpB2 and CdpB3 in wild type H26, *arcoA* L4*, *arcoB* A4*, Δ*arcoC*, and *arco* deletion strains. All GFP fusion proteins were expressed from plasmids under the control of their native promoters, respectively. Cells were grown at 45 °C overnight. Scale bars, 5 μm.

b-c. Representative images of the FtsZ2-GFP localization in wild type H26 or Δ*arcoABC* cells upon overexpression of individual Arco proteins. FtsZ2-GFP under control of its native promoter and Arco proteins under control by the $P_{tna}$ promoter are carried on the same plasmid. Cells were grown at 45 °C overnight with the indicated concentrations of tryptophan (Trp). Scale bars, 5 μm.

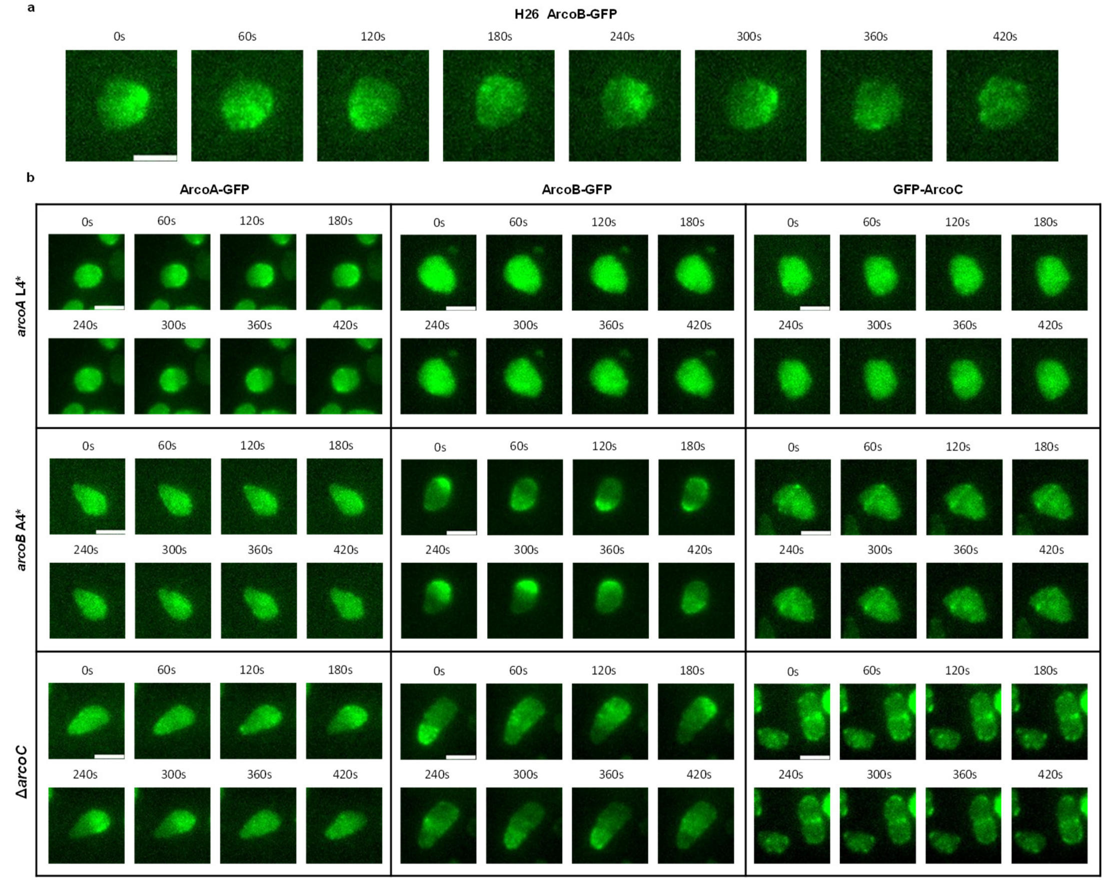


**Fig. S4 Oscillation modes and dependency of the Arco system.**

a. Representative time-lapse images of ArcoB-GFP in disc-shaped H26 cells. ArcoB-GFP was expressed from a plasmid under the control of the $P_{xyl}$ promoter, induced with 0.5 mM xylose at 45 °C overnight before imaging. Time in seconds.

b. Representative time-lapse images of ArcoA-GFP, ArcoB-GFP and GFP-ArcoC in *arcoA* L4*, *arcoB* A4*, and Δ*arcoC* strains. Oscillation dependency: ArcoA and ArcoB require each other and ArcoC requires both ArcoA and ArcoB. ArcoA-GFP was expressed from a plasmid under the control of the $P_{tna}$ promoter (induced with 1 mM Trp). ArcoB-GFP was expressed from a plasmid under the control of the $P_{xyl}$ promoter (induced with 0.5 mM xylose). GFP-ArcoC was expressed from a plasmid driven by the native promoter of the *arco* operon. All strains were grown at 45 °C overnight before imaging. Time in seconds.

Scale bars, 2 μm.

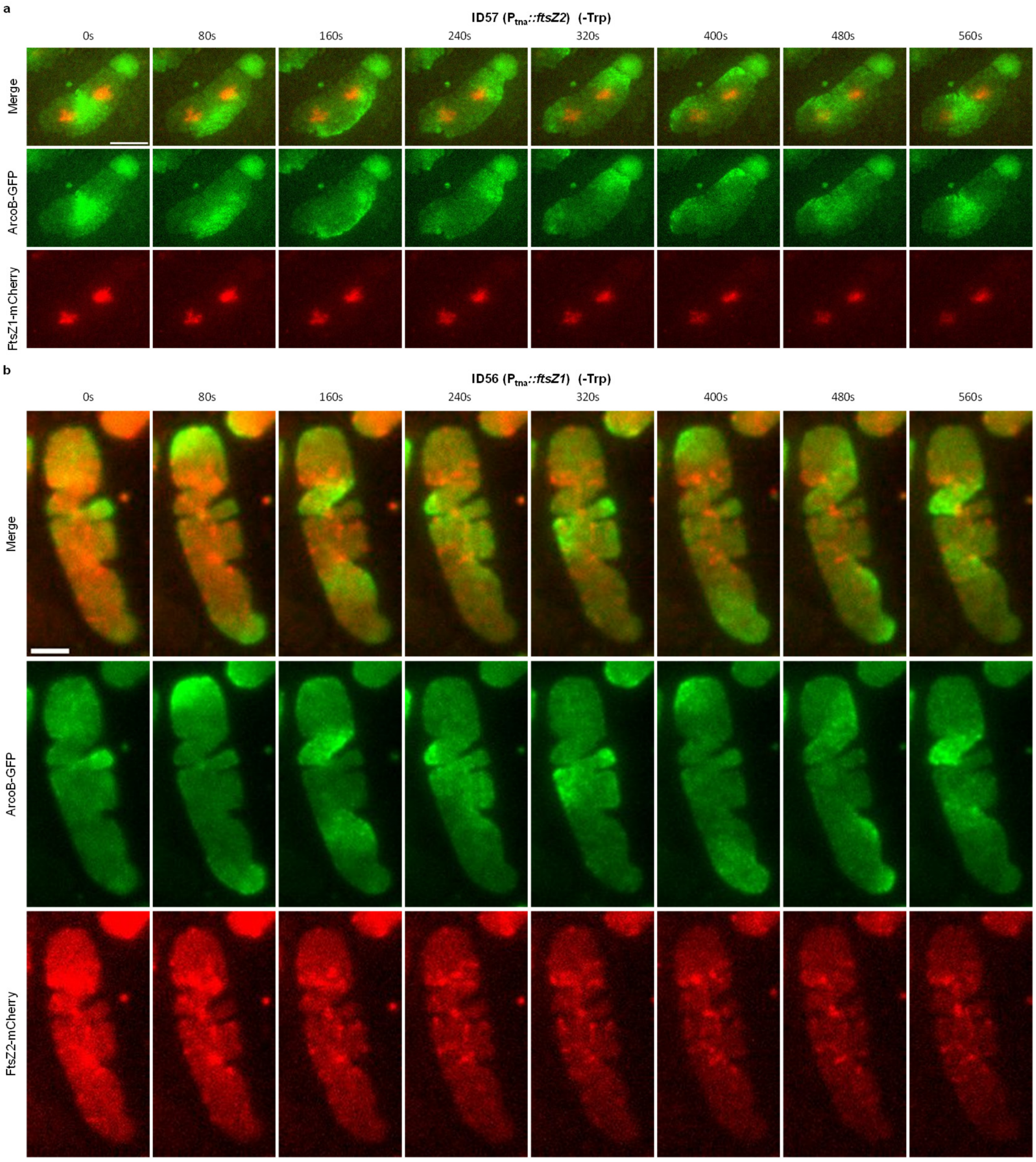


**Fig. S5 Arco oscillations restricts FtsZ1 localization but not FtsZ2.**

a. Representative time-lapse images of FtsZ1-mCherry and ArcoB-GFP in FtsZ2 depleted cells. Depletion of FtsZ2 was achieved by removal of tryptophan (Trp) from the cultures. FtsZ1-mCherry was under the control of its native promoter, and ArcoB-GFP was under the control of the $P_{xyl}$ promoter on the same plasmid. Cells were grown at 45 °C overnight with 0.5 mM xylose. Scale bar, 5 μm.

b. Representative time-lapse images FtsZ2-mCherry and ArcoB-GFP in FtsZ1 depleted cells. Depletion of FtsZ1 was achieved by removal of tryptophan (Trp) from the cultures. FtsZ1-mCherry was under the control of its native promoter, and ArcoB-GFP was under the control of the $P_{xyl}$ promoter on the same plasmid. Cells were grown at 45 °C overnight with 0.5 mM xylose. Scale bar, 2 μm.

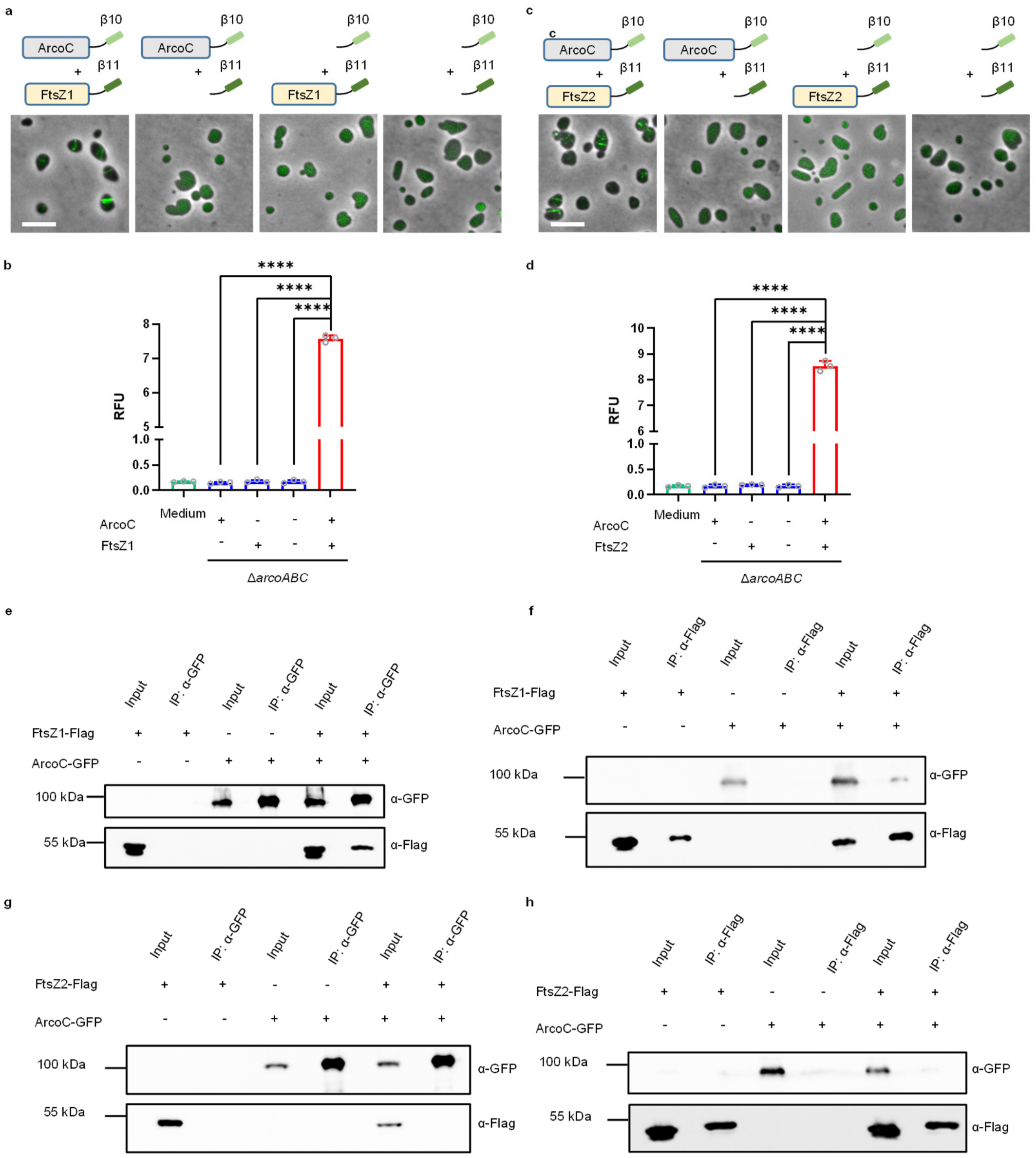


**Fig. S6 ArcoC interacts with FtsZ1 *in vivo*.**

a. Representative images of cells co-expressing ArcoC and FtsZ1 fused with the complementary fragments of superfolder GFP in the Split-FP assay.

b. Quantitation of the interaction signal between ArcoC and FtsZ1. RFU, relative fluorescence unit. Data are presented as mean values ± s.d. Significance in each group was tested by two-sided t-test. ****P <0.0001, n=3.

c. Representative images of cells co-expressing ArcoC and FtsZ2 fused with the complementary fragments of superfolder GFP in the Split-FP assay.

d. Quantitation of the interaction signal between ArcoC and FtsZ2. RFU, relative fluorescence unit. Data are presented as mean values ± s.d. Significance in each group was tested by two-sided t-test. ****P <0.0001, n=3.

e-h. Co-IP experiment showing that ArcoC interacts with FtsZ1 but not FtsZ2 *in vivo*. Cultures of *H. volcanii* expressing the indicated proteins were lysed by sonication; supernatants were incubated with anti-GFP (e, g) or anti-His (f, h) antibodies coated magnetic beads. Immunocomplexes were eluted with boiling SDS-PAGE loading buffer and then analyzed by immunoblot.

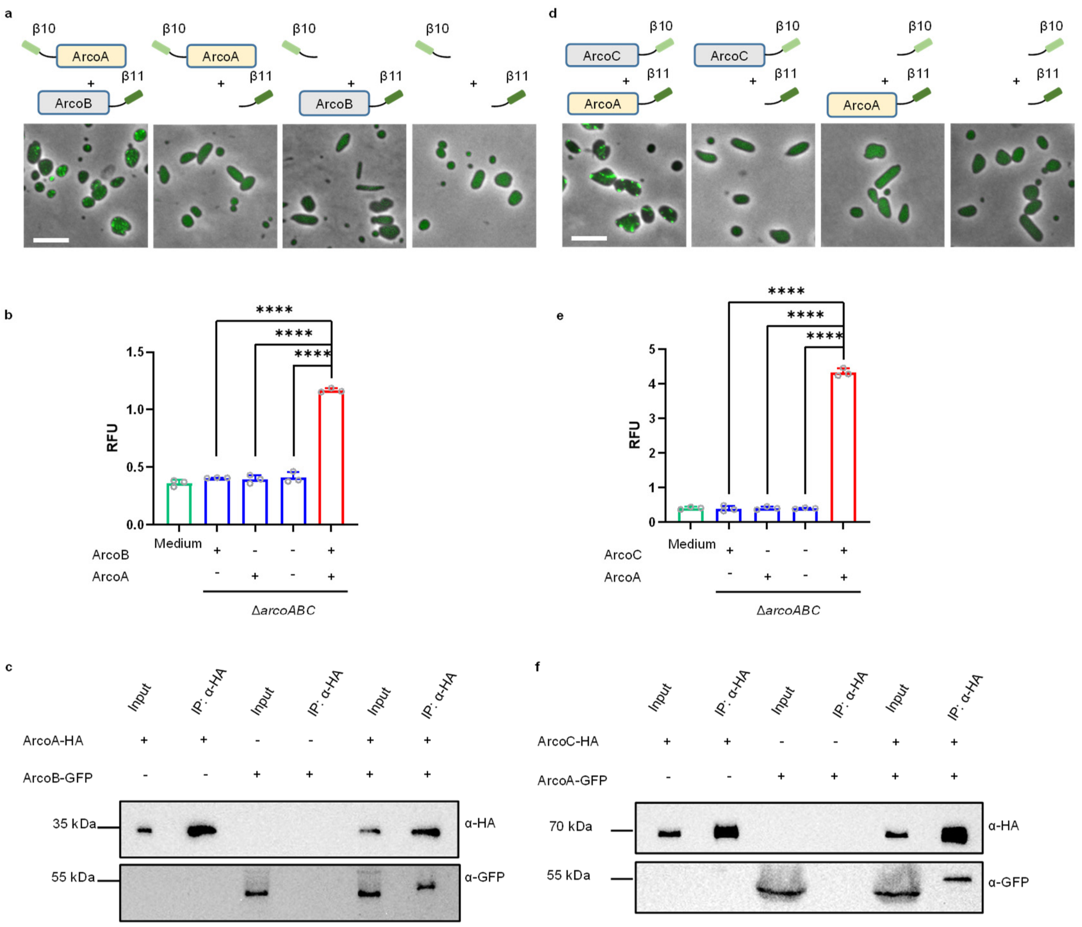


**Fig. S7 ArcoA interacts with both ArcoB and ArcoC *in vivo*.**

a. Representative images of cells co-expressing ArcoA and ArcoB fused with the complementary fragments of superfolder GFP in the Split-FP assay.

b. Quantitation of the interaction signal between ArcoA and ArcoB. RFU, relative fluorescence unit. Data are presented as mean values ± s.d. Significance in each group was tested by two-sided t-test. ****P <0.0001, n=3.

c. Representative images of cells co-expressing ArcoA and ArcoC fused with the complementary fragments of superfolder GFP in the Split-FP assay.

d. Quantitation of the interaction signal between ArcoA and ArcoC. RFU, relative fluorescence unit. Data are presented as mean values ± s.d. Significance in each group was tested by two-sided t-test. ****P <0.0001, n=3.

e-f. Co-IP experiment showing that ArcoA interacts with both ArcoB and ArcoC *in vivo*. Cultures of *H. volcanii* expressing the indicated proteins were lysed by sonication; supernatants were incubated with anti-HA antibodies coated magnetic beads. Immunocomplexes were eluted with boiling SDS-PAGE loading buffer and then analyzed by immunoblot.

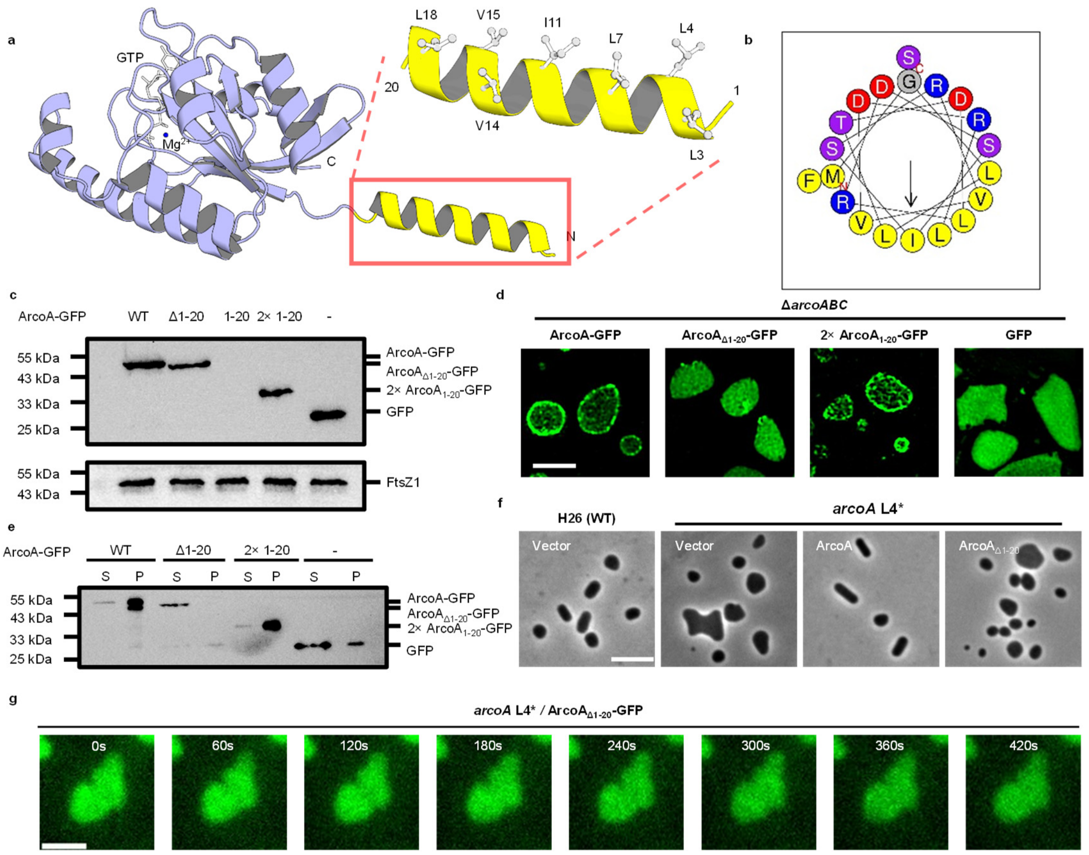


**Fig. S8 The amphipathic helix of ArcoA is essential for membrane binding and oscillation.**

a. The amphipathic helix of ArcoA revealed by Alphafold3 structure model. Atom colors: gray, carbon; blue, magnesium ion.

b. HeliQuest analysis of the amphipathic helix of ArcoA. Color code: yellow (non-polar), purple (polar), blue (charged), gray (Gly). Arrow points to the hydrophobic face.

c. Western blot analysis of the expression levels of ArcoA-GFP and its variants. GFP and GFP fusion proteins were all expressed from plasmids under the control of the $P_{tna}$ promoter, induced with 1 mM tryptophan (Trp) overnight at 45 °C. The cultures expressing the indicated proteins were normalized to the same $OD_{600}$, then lysed by sonication. The supernatants were mixed with 5× SDS-PAGE loading buffer, boiled for 10 min, and analyzed by immunoblotting. FtsZ1 serves as the loading control.

d. Representative images of the localization of ArcoA-GFP and its variants by 2D-SIM. GFP and GFP fusion proteins were expressed in Δ*arcoABC* cells from plasmids under the control of the $P_{tna}$ promoter, induced overnight with 1 mM tryptophan (Trp) at 30 °C. Scale bar, 2 μm.

e. Membrane fractionation assay showing that 2×ArcoA$_{1-20}$-GFP binds to the cell membrane, and the absence of amphipathic helix abolished the membrane-binding ability of ArcoA-GFP *in vivo*. Cultures of Δ*arcoABC* cells expressing the indicated proteins were lysed by sonication; the membrane fraction was isolated by ultracentrifugation from the supernatant; the pellet was resuspended to the original volume and analyzed by immunoblot. S, supernatant (cytosol); P, pellet (membrane).

f. Representative images showing that ArcoA$_{\Delta 1-20}$ cannot rescue the cell morphology and division defects of *arcoA* L4* cells. ArcoA or ArcoA$_{\Delta 1\text{-}20}$ was expressed from its native promoter on a plasmid. All strains were grown at 45 °C overnight before imaging. Scale bar, 5 μm.

g. Representative time-lapse images of ArcoA$_{\Delta 1\text{-}20}$-GFP in *arcoA* L4* cells. ArcoA$_{\Delta 1\text{-}20}$-GFP was expressed from a plasmid under the control of the $P_{tna}$ promoter (induced with 1 mM Trp). Cells were grown at 45 °C overnight before imaging. Time in seconds. Scale bar, 2 μm.

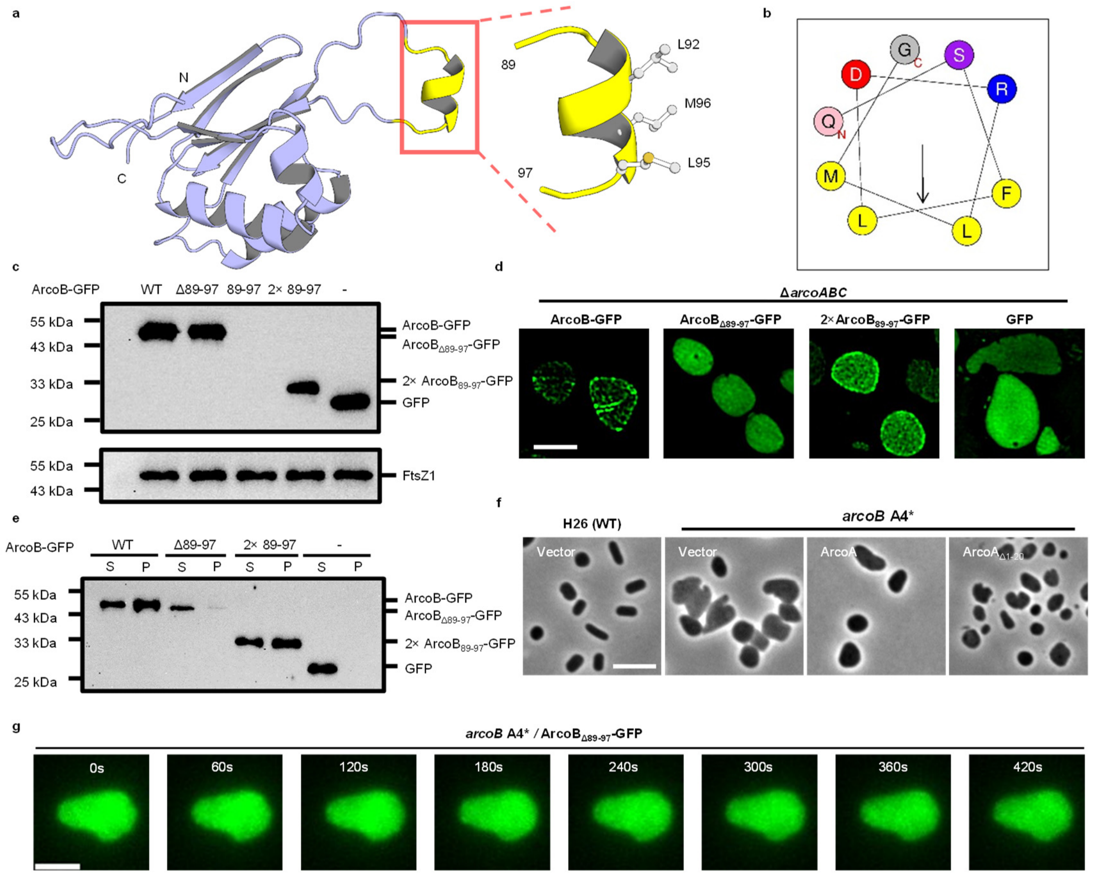


**Fig. S9 The amphipathic helix of ArcoB is essential for membrane binding and oscillation.**

a. The amphipathic helix of ArcoB revealed by Alphafold3 structure model. Atom colors: gray, carbon; orange, sulfur.

b. HeliQuest analysis of the amphipathic helix of ArcoB. Color code: yellow (non-polar), purple (polar), blue (charged), gray (Gly). Arrow points to the hydrophobic face.

c. Western blot analysis of the expression levels of ArcoB-GFP and its variants GFP-tagged proteins were all expressed from plasmids under the control of the $P_{xyl}$ promoter, induced with 0.5 mM xylose overnight at 45 °C. The cultures expressing the indicated proteins were normalized to the same $OD_{600}$, then lysed by sonication. The supernatants were mixed with 5× SDS-PAGE loading buffer, boiled for 10 min, and analyzed by immunoblotting. FtsZ1 serves as the loading control.

d. Representative images of the localization of ArcoB-GFP and its variants by 2D-SIM. GFP and GFP fusion proteins were expressed in Δ*arcoABC* cells from plasmids under the control of the $P_{xyl}$ promoter, induced overnight with 0.5 mM xylose at 30 °C. Scale bar, 2 μm.

e. Membrane fractionation assay showing that 2×ArcoB$_{89-97}$-GFP binds to the cell membrane, and the absence of amphipathic helix abolished the membrane-binding ability of ArcoB-GFP *in vivo*. Cultures of Δ*arcoABC* cells expressing the indicated proteins were lysed by sonication; the membrane fraction was isolated by ultracentrifugation from the supernatant; the pellet was resuspended to the original volume and analyzed by immunoblot. S, supernatant (cytosol); P, pellet (membrane).

f. Representative images showing that ArcoB$_{Δ89\text{-}97}$ cannot rescue the cell morphology and division defects of *arcoB* A4* cells. ArcoB or$_{Δ89\text{-}97}$ was expressed from the native promoter of the *arco* operon on a plasmid. All strains were grown at 45 °C overnight before imaging. Scale bar, 5 μm.

g. Representative time-lapse images of ArcoB$_{Δ89\text{-}97}$-GFP in *arcoB* A4* cells. ArcoB$_{Δ89\text{-}97}$-GFP was expressed from a plasmid under the control of the $P_{xyl}$ promoter (induced with 0.5 mM xylose). Cells were grown at 45 °C overnight before imaging. Time in seconds. Scale bar, 2 μm.
